# A thalamic circuit represents dose-like responses induced by nicotine-related beliefs in human smokers

**DOI:** 10.1101/2022.07.15.500226

**Authors:** Ofer Perl, Anastasia Shuster, Matthew Heflin, Soojung Na, Ambereen Kidwai, Natalie Booker, William C. Putnam, Vincenzo G. Fiore, Xiaosi Gu

## Abstract

Could non-pharmacological constructs, such as beliefs, impact brain activities in a dose-dependent manner as drugs do? While beliefs shape many aspects of our behavior and wellbeing, the precise mapping between subjective beliefs and neural substrates remains elusive. Here, nicotine-addicted humans were instructed to think that an electronic cigarette (e-cigarette) contained either “low”, “medium”, or “high” levels of nicotine, while nicotine content was kept constant. After vaping the e-cigarette, participants performed a decision-making task known to engage neural circuits affected by nicotine while being scanned by fMRI. Activity in the thalamus, a key binding site for nicotine, increased parametrically according to belief dosage. Furthermore, the functional coupling between thalamus and ventromedial prefrontal cortex, a region implicated in value and state representations, also scaled to belief dosage. These findings illustrate a dose-dependent relationship between a thalamic circuit and nicotine-related beliefs in humans, a mechanism previously known to only apply to pharmacological agents.

## Introduction

Humans hold beliefs that can profoundly alter our behaviors and wellbeing. Albeit subjective in nature, beliefs – similar to other mental functions – are represented by biological substrates in the brain^1,2^. However, the exact mapping between subjective beliefs and neurobiological substrates remains largely unknown, hindering our understanding of neuropsychiatric conditions like drug addiction, where purely biochemical explanations are not sufficient to account for the complexity of the disorder^3,4^. Elucidating the precise neural mechanisms of beliefs is also important for understanding how beliefs and expectations play a role in pharmacological treatments, where individuals’ drug response differ drastically^5^.

The placebo effect represents a notable example supporting the potential interaction between beliefs and neurobiology. In these observations, one’s symptoms can improve simply due to positive beliefs about a receiving a treatment while there is no active ingredient in the drug^5–9^. Even in the presence of a powerful neuroactive substance such as nicotine, beliefs can exert a binary all-or-none type of effect on neural responses in human smokers. Collectively, these findings provide initial support for the notion that beliefs can broadly affect neurobiological activities in the human brain^10,11^ without evidence for how precise the neural effect of beliefs might be.

To establish precision, pharmacological research has typically relied on the concept of dose-dependent response, where the amount of active ingredients in a drug is known to modulate biological processes proportionally. In terms of neuropharmacology, dose responses in the brain have been observed in a wide range of neuroactive drugs such as nicotine^12^, alcohol^13^, and marijuana^14^. However, such inquiry has rarely existed in neuroscience research on human beliefs. Is it possible that beliefs – a highly subjective and implicit mental construct – could modulate neurophysiological responses in a similar dose-dependent manner?

Based on the literature reviewed thus far, we hypothesized that human beliefs – such as those related to neuroactive substances like nicotine – can modulate brain activities in a manner that is similar to pharmacologically induced dose responses. Nicotine is known to broadly affect distributed regions in the brain, including the thalamus and the striatum^15,16^, both of which are important for cognition and decision-making^17,18^. The thalamus in particular, contains one of the highest densities of nicotinic acetylcholine receptors (nAChRs) for nicotine binding in the human brain^19,20^. The stimulation of nAChRs by nicotine can lead to subsequent dopamine release in mesolimbic structures such as the ventral striatum^15,21^. In humans, however, high levels of nicotine are not a necessary condition for the activation of nAChRs. For instance, positron emission tomography (PET) imaging studies have demonstrated that there can be a substantial degree of occupancy of nAChRs even when nicotine-addicted individuals only smoked denicotinized or low-nicotine content cigarettes^22,23^, or only had second-hand smoke^24^. These findings pinpoint to the possibility that nicotine itself was not sufficient to account for the complex neural effects observed in nicotine-dependent humans, and that cognitive constructs such as nicotine-related beliefs may play a crucial role in modulating addiction neurobiology.

To test this hypothesis, we instructed nicotine-dependent human smokers to believe that an electronic cigarette (e-cigarette) they were about to vape contained either “low”, “medium”, or “high” levels of nicotine, while the actual nicotine content was fixed across all e-cigarettes (see **Materials and Methods** for details). After vaping, smokers (final sample included 60 scans across 20 smokers) performed a monetary decision-making task during functional magnetic resonance imaging (fMRI; **Fig. 1a**). A group of non-smoking healthy controls (HCs, n=31) also performed the same fMRI task, but without going through the vaping procedure. We chose to use e-cigarettes to deliver nicotine as nicotine strength can be controlled much more precisely compared to traditional cigarettes. Based on the literature reviewed thus far, we predicted that the activity in those neural regions characterized by high nAChRs (i.e. thalamus and related structures) might represent beliefs about nicotine dose in a precise manner, resembling the dose-dependent responses found in pharmacological studies. If proven true, such a finding would reveal a higher degree of sophistication and precision of the mapping between human beliefs and brain states than previously understood.

**Fig. 1.**
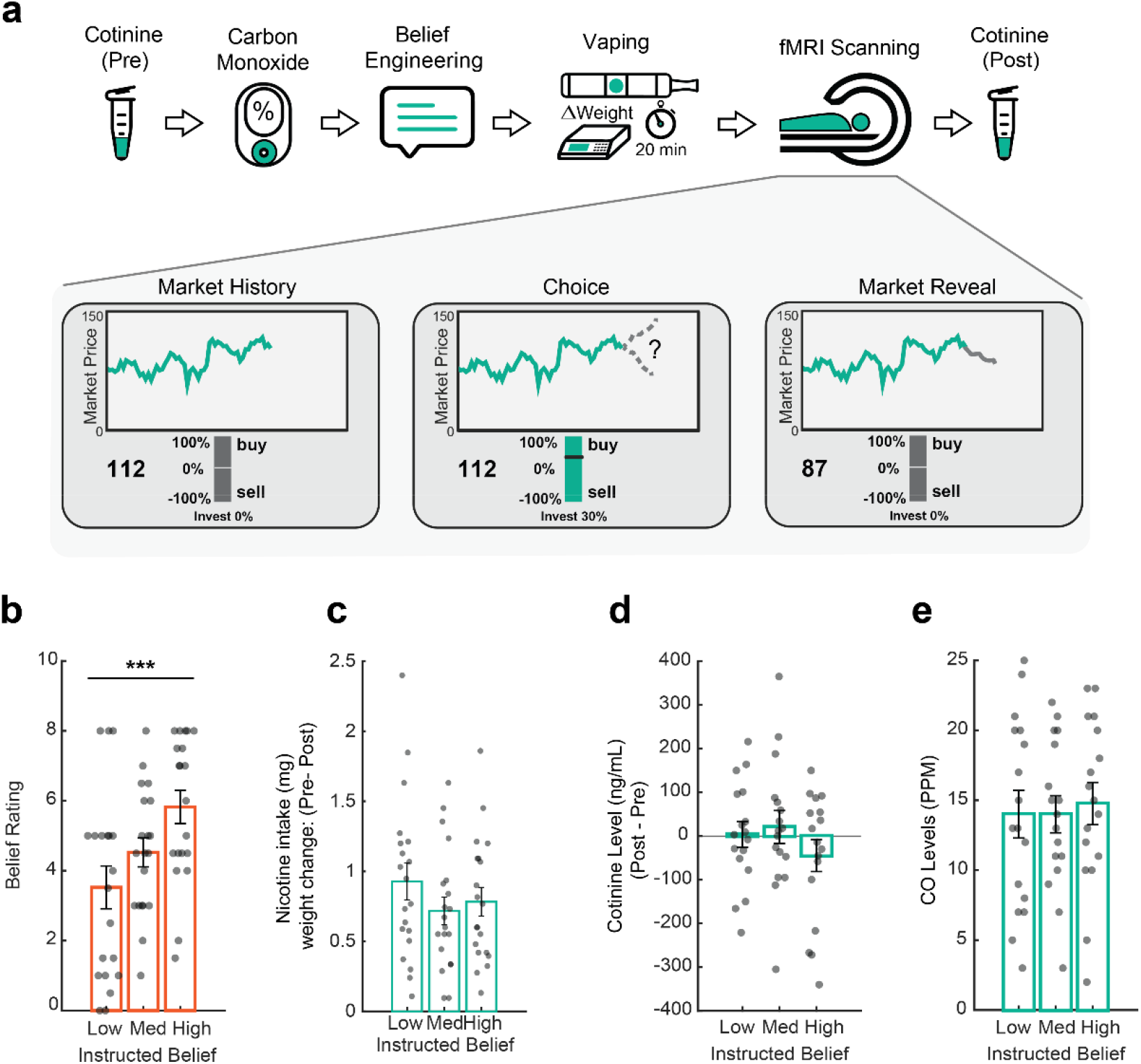
Experimental paradigm and sanity check measures. (**a**) Participants completed three visits. In each visit, we collected saliva samples for cotinine measurement, measured carbon monoxide (CO) levels, instructed beliefs, and measured brain activities using fMRI as participants engaged in a decision-making task. (**b**) Subjective beliefs about nicotine strength increased as a function of instructed nicotine strength (*P* = 0.0004). (**c**) Consumed nicotine was similar across three belief conditions (*P* = 0.178), (**d**) cotinine concentration (*P* = 0.393), or (**e**) CO level (*P* = 0.698) did not differ between conditions. Bars depict group means and points represent participants. Error bars are SEM. \*\*\**P* < 0.001.

## Results

### Instructions induced changes in subjective beliefs in smokers, but did not affect overall nicotine intake, metabolism, or baseline saturation

Our key experimental manipulation is the instruction given to the participants about whether the e-cigarette had “low”, “medium”, or “high” strength of nicotine. To examine if this design indeed induced changes in beliefs about nicotine in our subjects, we asked all participants to rate their perceived nicotine strength using a 10-point scale after vaping. Overall, participants’ perceived nicotine strength significantly increased as a function of instructed beliefs about nicotine dosage (mean ± SD (AU); ‘low’: = 3.52 ± 0.61, ‘medium’: = 4.52 ± 0.41, ‘high’: = 5.82 ± 0.47, rmANOVA *F*(2,38) = 9.71, *P* = 0.0004, partial η2□=□0.34, 90% CI□=□0.12≤□η□≤□048, **Fig. 1b**), supporting the validity of our experimental manipulation.

Next, we took extensive sanity checks to ensure the instruction did not interfere with participants’ nicotine intake, metabolism, or their baseline nicotine saturation levels. First, participants might vape less in “weaker” nicotine conditions due to a lack of interest. To control for this, we set vaping time to 20 minutes during data collection. Importantly, we also quantified the amount of nicotine intake, which equals the change in cartridge weight after vaping multiplied by the actual percentage of nicotine content (1.2%). We found that nicotine intake did not differ across belief conditions (nicotine intake (mg): ‘low’: = 0.928 ± 0.56, ‘medium’: = 0.719 ± 0.423, ‘high’: = 0.783 ± 0.434; rmANOVA *F*(2,38) = 1.806, *P* = 0.178; **Fig. 1c**), suggesting that difference in belief about nicotine did not affect how much liquid or nicotine was consumed by the smokers. The overall amount of consumed nicotine here is in a range that is similar to nicotine delivered by traditional cigarettes in previous experimental studies^10,11,23^.

However, it might be possible that even with the same nicotine consumption, nicotine metabolism might still differ between belief conditions. To address this, we collected saliva samples both before and after vaping for high-performance liquid chromatography tandem mass spectrometry (LC-MS/MS) analytical quantification of cotinine—a nicotine metabolite indicative of plasma nicotine levels^25^ (see **Materials and Methods** for details). We found that vaping-induced changes in cotinine concentrations (ng/mL) were comparable across conditions (rmANOVA *F*(2,32) = 0.959, *P* = 0.393; **Fig. 1d)**, suggesting that nicotine metabolism itself was unlikely a factor contributing to any brain-based differences.

We also measured exhaled carbon monoxide (CO) before vaping as an index of participants’ baseline nicotine saturation level. We did not observe differences in CO levels across conditions (parts per million (ppm); rmANOVA *F*(2,32) = 0.364, *P* = 0.698; **Fig. 1e**). Taken together, these analyses confirmed that our instruction successfully influenced participants’ beliefs about nicotine strength, while mitigating the concern that imbalanced nicotine consumption, metabolism, or baseline deprivation might have contributed to any neural differences across conditions.

### Thalamic representation of dose-like responses induced by nicotine-related beliefs

Our main quest here is how beliefs about nicotine are represented by neural activities in smokers. We chose to measure neural activities during a value-based decision-making task because both nicotine and belief about nicotine have been shown to influence similar circuitries involved in reward processing^10,11,26,27^. Specifically, we used a sequential investment task (see **Materials and Methods** for details) to probe reward processing; similar paradigms have been previously used in both healthy controls and those with nicotine addiction^10,11,26,27^. Briefly, participants made a series of choices regarding how to invest (or short-sell) in simulated stock markets, based on one’s prediction of market return *r*_*t*_, defined as *r*_*t*_ *= (p*_*t*_*−p*_*t−1*_*) / p*_*t−1*_ (where *p*_*t*_ denotes the market price at time *t*). Because subjects were allowed to place either positive or negative bets, they could win (or lose) money in either positive or negative markets. As such, the absolute value |*r*| represents the actual reward value that is attainable to the subject.

In a whole-brain ANOVA with belief as the main factor (“low”, “medium”, or “high”) and the value signal |*r*_*t*_| as the key parametric modulator (see **Materials and Methods** for details), we observed that value-related neural activities in the thalamus exhibited a dose-dependent response to instructed beliefs about nicotine strength (peak at MNI: x = -15, y = -19, z = -1; *P* < 0.05, FWE (family-wise error) cluster-corrected at a cluster-defining threshold of *P* < 0.005, uncorrected, *k* = 50, *P* = 0.006; **Fig. 2a**). No other brain structures showed a similar neural activity pattern in relation to beliefs at the whole brain level with the same statistical threshold.

**Fig. 2.**
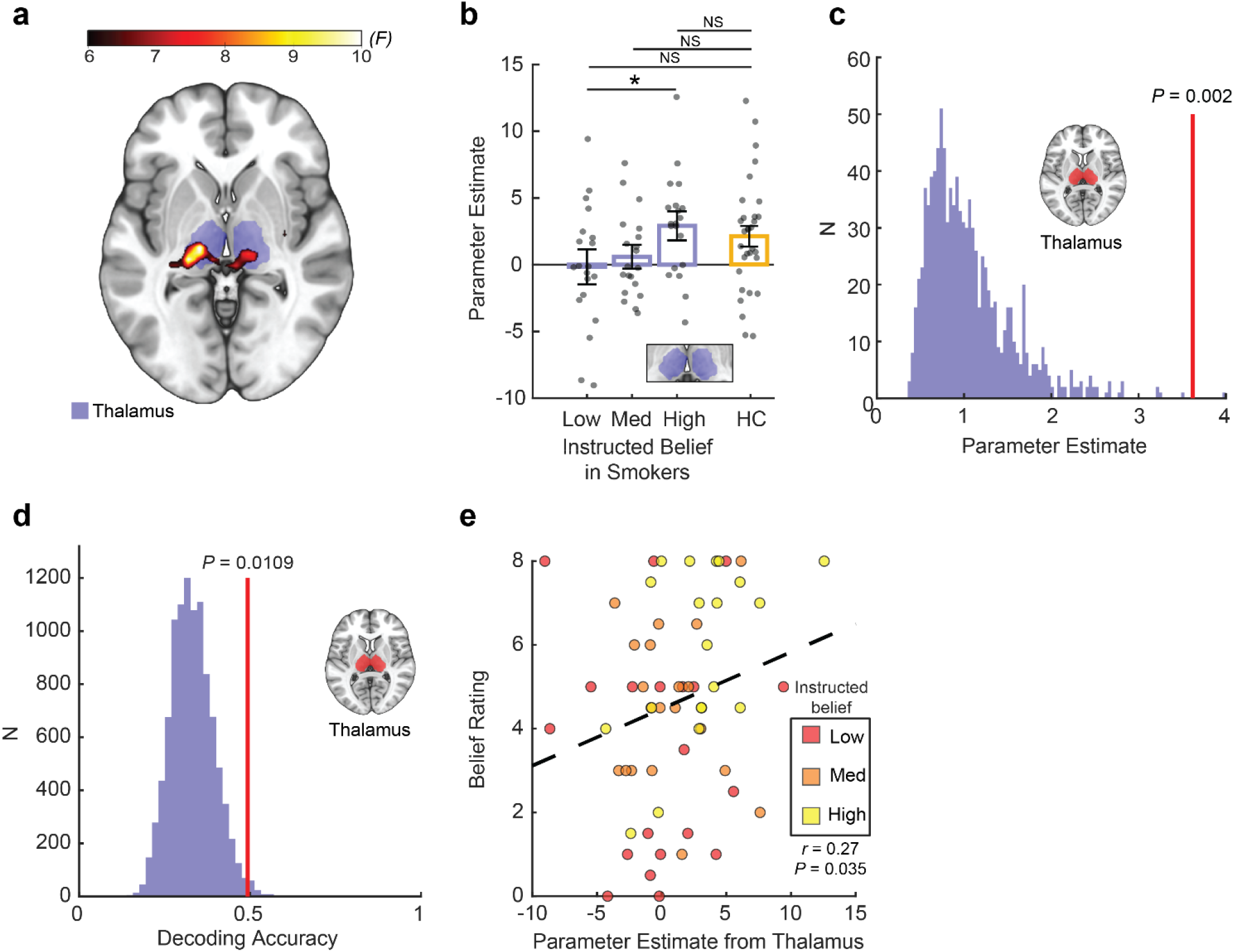
Belief about nicotine strength induced dose-dependent responses in the thalamus. **(a)** Whole-brain effects of instructed beliefs about nicotine on value-tracking signals (rmANOVA, cluster-level *P*_*FWE*_ = 0.006, *k* = 50). **(b**) Parameter estimates representing reward-related activities extracted from an independent thalamus mask (shown in purple) across belief conditions in smokers (*P* = 0.036), compared to a non-smoking healthy controls (HC, orange bar) Bars depict group means, points represent participants. Error bars are SEM. \**P* < 0.05. (**c**) Permutation analysis for instructed beliefs conditions (*N* = 1,000, *P* = 0.002). A histogram comprised of surrogate distribution for beta estimates (black bars). Red line denotes mean of true beta values. **(d)** Decoding accuracy of belief condition from thalamic neural patterns. Vertical red line denotes decoding accuracy for ground truth data. Colored histogram is a surrogate distribution comprised of decoding accuracy for the same neural data with shuffled labels. P value is derived non-parametrically through a permutation test (*N* = 10,000). (**e**) Correlation between thalamic signals and subjective belief rating regarding perceived nicotine strength (r = 0.27, *P* = 0.035). Black dashed line is linear fit.

Whole brain-level statistical maps of each belief condition are available in **Supplementary Fig. 1)**.

A region of interest (ROI) analysis using an independent anatomical mask^28^ further confirmed that BOLD signals from the thalamus differentiated between instructed belief conditions (mean ± SD (AU); ‘low’: = 0.157 ± 1.047, ‘medium’: = 0.601 ± 0.714, ‘high’: = 2.914 ± 0.865; rmANOVA test *F*(2,38) = 3.62, *P* = 0.036, partial η2□=□0.16, 90% CI□=□0.0057□≤□η□≤□0.30, **Fig. 2b)**. These activations did not differ significantly from non-smoking health controls (n=31 for HCs; see **Materials and Methods** for details; mean ± SD (AU) for HC: = 2.318 ± 4.258; two-sample t-test: ‘smokers-low’ vs ‘HC’ t(49) = 1.95, p = 0.057; ‘smokers-medium’ vs ‘HC’ t(49) = 1.53, p = 0.132; ‘smokers-high’ vs ‘HC’ t(49) = -0.50, p = 0.616).

We also carried out a non-parametric approach to examine if the relationship we observed between beliefs and neural activities was indeed not random. Using a permutation analysis approach, we iteratively extracted beta estimates from surrogate GLMs based on shuffled belief conditions (*N* = 2,000). We observed that beta estimates for the actual allocation of belief conditions ranked significantly higher than the surrogate distribution (*P* = 0.002, **Fig. 2c)**. A finer parcellation of the thalamus^29^ revealed that ventral posterior nuclei – notably the centromedian (CM) and lateral geniculate nuclei (LGN), were the primary nuclei in which reward-tracking neural activity differentiated between instructed beliefs in a parametric manner (FDR corrected at q = 0.05; VPL, Pulvinar, LGN, CM all P < 0.05; see **Supplementary Information** and **Supplementary Fig. 2**).

Next, we asked whether thalamic activities were actually predictive of the belief condition using a decoding analysis. We trained a regularized linear discriminant analysis (rLDA) model to decode instructed belief conditions from multivoxel spatial patterns extracted from the thalamus^30,31^. We were able to decode at 49.3 % accuracy the instructed belief condition from the distributed multivoxel patterns of thalamic activity. This decoding accuracy was significantly greater than chance level (33.3 %), as confirmed by a permutation test where we iterated the procedure with shuffled labels (N = 10,000) and compared the true decoding accuracy to the surrogate accuracy distribution (surrogate: 33.1 ± 6.3 %, *P* = 0.011, **Fig. 2d**). We further applied this decoding approach to each nucleus within the thalamus, using the same anatomical parcellation as before. We observed that decoding accuracy was roughly aligned with the spatial distribution of effects uncovered in the GLM analysis in that there was greater decoding accuracy in the ventral posterior nuclei. Following FDR correction only the posterolateral nucleus (VPL) nucleus showed decoding ability significantly higher than chance (FDR corrected at q = 0.05; VPL, P = 0.018; see **Supplementary Fig**. 4).

Given the individual variability in susceptibility to instructed beliefs^10,11^, we also asked whether participants’ subjective beliefs, indexed by their self-reported perception about nicotine strength, also parametrically modulated thalamic responses. We found that across all participants and all sessions, subjective ratings of perceived nicotine strength correlated with reward-related activities in the thalamus (Spearman correlation, *r* = 0.27, *P* = 0.035, **Fig. 2e**), suggesting that these neural signals were linked to participants’ perceptions about nicotine strength following instructed beliefs. Taken together, these analyses further confirmed that experimental instructions about nicotine strength shaped subjective perception in smokers and induced dose-dependent neural responses in the thalamus, a brain region with one of the highest concentrations of nicotinic acetylcholine receptors and a main binding site for nicotine^19,32^.

Taken together, these results pinpoint to the thalamus – in particular the posterior thalamus - as a key neural substrate representing nicotine-related beliefs. This finding might provide a mechanistic account for the previously observed effects that smoking denicotinized or low-nicotine content cigarettes can still induce a substantial level of nAChR occupancy in the human brain^22–24^.

### Observed effect of beliefs on thalamic activity was not due to sensorimotor effects or spatial smoothing

Next, we conducted several control analyses to rule out alternative explanations of observed thalamic effects. In addition to its involvement in nicotine addiction, the thalamus - especially the thalamic nuclei identified so far - is also known for encoding basic sensorimotor information. Thus, it is possible that the differential neural states in the thalamus were induced by different levels of visual or motor processing for the three conditions, instead of the belief per se. To rule out this possibility, we first checked button pressing behavior during the task, and found no difference between belief conditions (see **Supplementary Information)**. We also examined neural responses related to button presses (motor) and simple viewing of market (visual) by constructing separate GLMs to model the fMRI data. We found no difference in thalamic activity related to motor or visual processing between belief conditions **(Supplementary Fig. 3a)**.

Given that technical choices during fMRI preprocessing such as spatial smoothing could have an impact on the resulting findings^33^, we also conducted all of our main analyses again by using a preprocessing pipeline without spatial smoothing (see Materials and Methods for details). We confirmed that the identified belief representation in thalamus was not due to spatial smoothing (**Supplementary Fig. 3b**). Taken together, these additional analyses ruled out several important confounds and suggest that it is unlikely that visual or sensorimotor elements contributed to the observed mapping between belief conditions and thalamic activity.

### Striatal activity tracked reward value, but did not distinguish between belief conditions

Thus far, our primary finding related on the key belief effect centered around the thalamus. However, previous work^10^ has also identified the ventral striatum as a key region that could be modulated by belief about nicotine. The ventral striatum, a mesolimbic region receiving dopaminergic inputs from the ventral tegmental area (VTA), is known to encode reward signals and also affected by nicotine addiction. Thus, we conducted a separate set of analyses focused on the ventral striatum. Consistent with previous findings, we found that the ventral striatum tracked the market value signal |*r*_*t*_| across all conditions (*P*_*FDR*_ q < 0.01, **Fig. 3a**). However, striatal responses did not differ between belief conditions at the whole brain level in an ANOVA analysis (P > 0.05, **Supplementary Fig. 5a**).

**Fig. 3.**
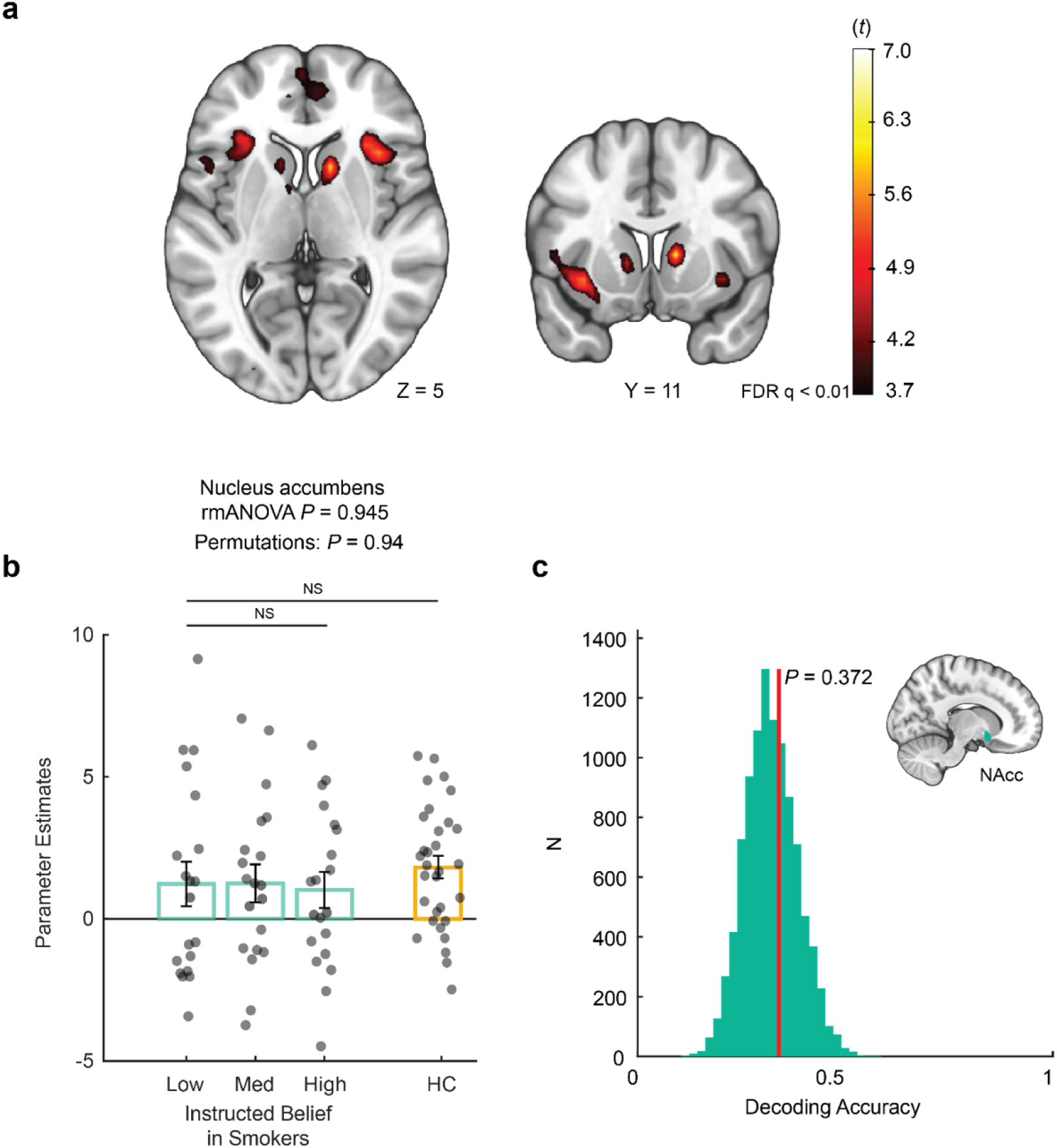
Belief about nicotine strength did not modulate striatal reward-related responses. **(a)** Whole-brain effects of cross-condition brain activation tracking market return across all instructed belief conditions. Heatmap signifies *t* values. **(b)** Parameter estimates representing reward-related activities extracted from an independent nucleus accumbens mask across belief conditions in smokers (teal bars) (rmANOVA *P* = 0.945; Permutations *P* = 0.94), compared to a non-smoking healthy controls (HC, orange bar). Bars depict group means, points represent participants. Error bars are SEM. **(c)** Decoding accuracy of belief condition from accumbens neural patterns. Vertical red line denotes decoding accuracy for ground truth data. Colored histogram is a surrogate distribution comprised of decoding accuracy for the same neural data with shuffled labels. P value is derived non-parametrically through a permutation test (*N* = 10,000).

**Fig. 4.**
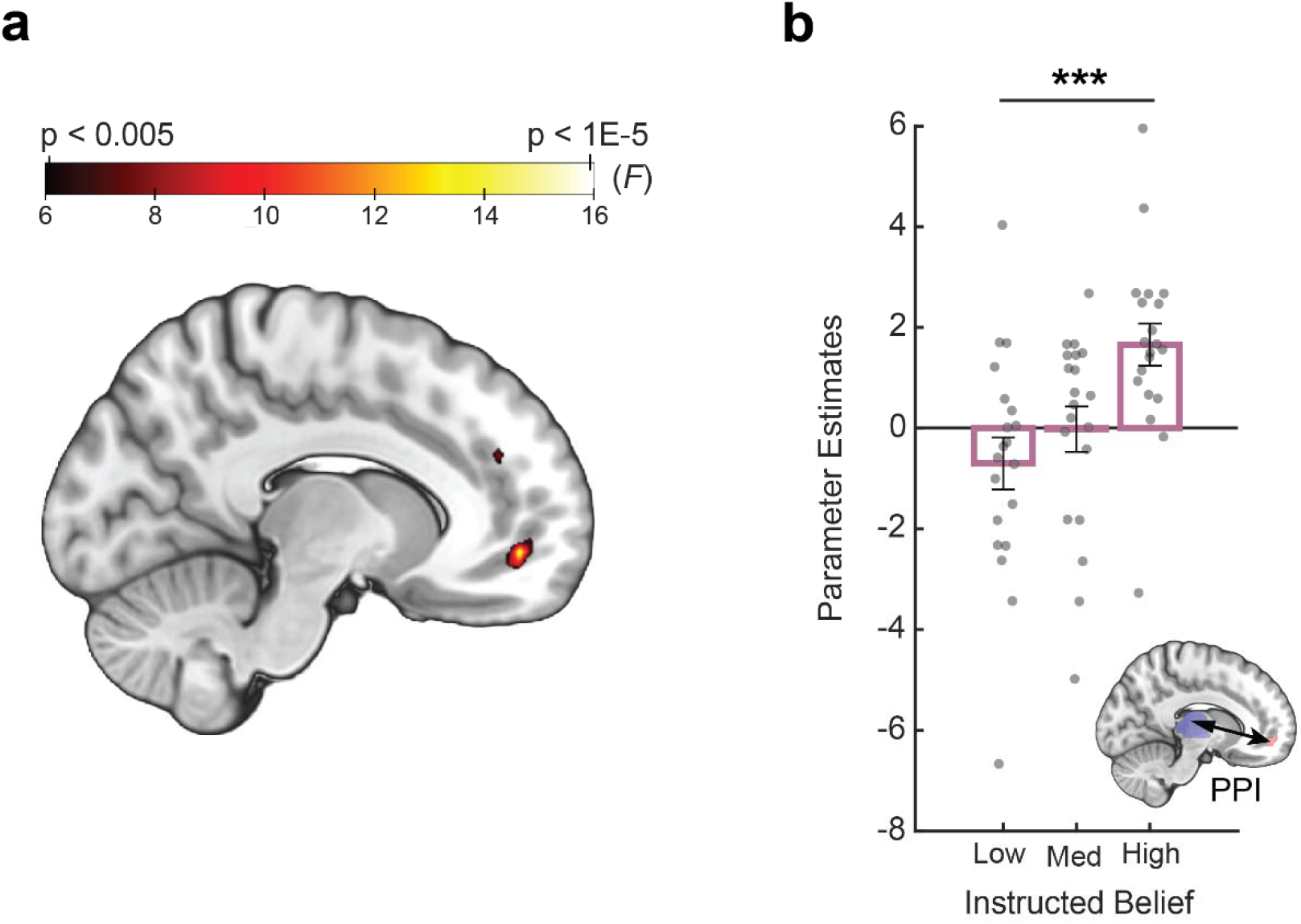
Belief about nicotine strength modulated thalamus-vmPFC functional connectivity in a dose-dependent fashion. (**a**) Effects of instructed beliefs on the psychophysiological interaction (PPI) between the thalamus and the vmPFC. (**b**) Parameter estimates extracted from (a) representing functional coupling strength between the thalamus and vmPFC. Bars depict group means, points represent participants. Error bars are SEM.

An ROI analysis using an independent mask of the nucleus accumbens (NAcc) further confirmed that neural activities in the NAcc did not differentiate between belief conditions (rmANOVA *F*(2,38) = 0.056, *P* = 0.945, permutation test: *P* = 0.94 ; **Fig. 3b**). These parameter estimates were also comparable to those extracted from the same group of healthy controls using the same NAcc mask (smokers: ‘low’: = 1.228 ± 3.329, ‘medium = 1.248 ± 2.828 ‘high’: = 1.016 ± 2.6983, ‘HC’: = 1.781 ± 2.138; two-sample t-test: ‘low’ vs ‘HC’ t(49) = -0.740, p = 0.462; ‘medium’ vs ‘HC’ t(49) = -0.784, p = 0.436; ‘high’ vs ‘HC’ t(49) = -1.138, p = 0.261).

In line with the GLM results, classification accuracy for belief condition using patterns extracted from the NAcc was not significantly higher than chance (ground truth = 34.0 %, surrogate: 32.1 ± 6.3 %, P = 0.372; **Fig. 3c**). Finally, we examined reward-related activations in other basal ganglia nuclei, namely the putamen and the caudate nucleus. We did not find significant differences between belief conditions either in separate ROI analyses (putamen rmANOVA *F*(2,38) = 1.15, *P* = 0.327; caudate rmANOVA *F*(2,38) = 0.24, *P* = 0.781; **Supplementary Fig. 5b-c**).

Seemingly surprising at a first glance, the lack of belief effects on the striatum was consistent with the lack of belief effect of instructed beliefs on reinforcement learning behavior in smokers in this study (see **Supplementary Information** and **Supplementary Fig. 6 and Supplementary Table 1** for details). Combined with the main belief effect concerning the thalamus, we speculate that the experimentally manipulated beliefs in this study primarily modulated low-level information gating as opposed to high-level value-guided decision-making in the previous study^10^. This difference might be attributed to the fact that smokers were not familiar with e-cigarettes in the current study and thus were not driven by conditioned responses tied to using a traditional cigarette as is the case for previous work^10^. We will discuss this in more detail later.

### Belief about nicotine modulated functional connectivity between prefrontal cortex and thalamus in a dose-dependent manner

At the circuit level, the thalamus is heavily connected to various cortical regions and is known to contribute to higher-order cognition via these connections^34^. Thus, we hypothesized that belief might also modulate the functional connectivity between the thalamus and higher cortical regions such as prefrontal regions. Specifically, the ventromedial prefrontal cortex (vmPFC) has been increasingly recognized as a key region in representing task states^35,36^ and the structure of abstract knowledge. Anatomically, it is well known that the thalamus and vmPFC are densely connected ^37,38^. Thus, we predicted that thalamic-vmPFC coupling would differ between belief conditions in our study.

To this end, we carried out a psychophysiological interaction (PPI; see **Materials and Methods**) analysis^39^ with the thalamus as a seed region to investigate how beliefs about nicotine were represented at a neural circuitry level. We found that belief about nicotine indeed modulated functional connectivity between the thalamus and the vmPFC both at the whole brain level (*P*_SVC_ < 0.05, FWE cluster-corrected at a cluster-defining threshold of *P* < 0.005, uncorrected, *P* = 0.041; **Fig. 3a**) and via an ROI analysis using a vmPFC mask from an independent study involving belief formation^40^ (peak at MNI x = -11, y= 50, z = -6, *k* = 5; **Supplementary Fig. 7a**; **Fig. 3b**). In sharp contrast, a separate set of PPI analyses using the ventral striatum as a seed region did not yield any significant changes in functional connectivity with the vmPFC or any other brain region at the same threshold (**Supplementary Fig. 7b)**. The vmPFC is a brain region heavily implicated in the computation of value and belief updating^36^ Importantly, recent work has pinpointed to the vmPFC for its representation of task states^41^. Thus, this result suggests that in addition to modulating thalamic activation itself, belief about nicotine also parametrically scaled circuit-level interactions between the thalamus and a prefrontal region involved in higher-level cognition and decision-making.

## Discussion

How are drug-related beliefs represented in the human brain? Using nicotine as a test case, we demonstrated that verbal instruction regarding nicotine strength (“low”, “medium”, or “high”) modulated how human smokers perceived the strength of nicotine contained in an e-cigarette that they vaped. Importantly, beliefs about nicotine strength were represented by neural activities in the thalamus in a dose-dependent fashion, during value-based decision-making. Across individuals, the subjective perception of nicotine strength parametrically correlated with neural activities in thalamus. At the circuitry level, the functional coupling between thalamus and vmPFC also scaled parametrically to belief “dose”. Taken together, these findings demonstrate the precise mapping between beliefs and neural activities in a prefrontal-thalamic circuit.

While humans hold beliefs about a wide range of stimuli and events, beliefs about substances are particularly important to examine due to their high relevance regarding substance use disorders. Here we demonstrated that nicotine-related beliefs are mapped onto neural states of the brain circuits that are critically involved in nicotine addiction in a way that mimics dose-responses of pharmacological agents. The thalamus — especially its posterior portion — contains one of the highest densities of nAChRs in the human brain^20^ as quantified by both autoradiography^19,42,43^ and functional imaging^44,45^. This anatomical feature is hypothesized to account for the attention-enhancing effect of nicotine^44,46^ as the thalamus is known to be central for gate incoming sensory information. Indeed, previous work has demonstrated acute dose-dependent responses induced by nicotine itself in the human thalamus^45^. Importantly, previous work showed that even when nicotine level was moderate or close to none, smoking cigarettes can induce a substantial level of occupancy of nAChRs in the thalamus in human smokers^22–24^. This suggested that habitual behaviors that had been previously reinforced by the intake of nicotine (e.g., the act of smoking itself) can modulate thalamic activity irrespective of actual nicotinic content. However, the mechanism linking the effect of this observable behavior to subjective states remained unclear. Our study further reveals a granular mechanism that might account for these previous findings – that difference in neural activations can be triggered by manipulating one’s beliefs about nicotine intake (which likely acts as precursors of explicit habitual actions), as if the nAChRs receptors were activated by the presence of actual different dosages of nicotine. This implies that cognitive constructs such as beliefs and expectations can modulate fine-grained biological mechanisms in the human brain in a way that is similar to pharmacological agents.

We also found that vmPFC-thalamus functional coupling during decision-making also distinguished between belief conditions. The vmPFC has been extensively studied in the context of value-based decision-making processes and has been proposed to encode a “common currency” of subjective value^47^. In serving this role, the vmPFC has been shown to receive input from both the ventral tegmental area and the basal ganglia via the thalamus^17^. Importantly, more recent computational accounts suggest that the vmPFC encodes task states, including forming abstract representations of task structures that are not directly observable^48^. Consistent with our current finding, the functional connectivity between vmPFC and thalamus has also been shown to subserve prior expectations about incoming visual stimuli^49^. Here, our finding expands previous work by demonstrating that instead of functioning as a binary “switch”, the vmPFC-thalamus circuit encodes information related to beliefs and expectations in a parametric manner, highlighting the importance and precision of this circuitry in representing abstract mental states.

In contrast to the thalamus, the ventral striatum tracked reward value overall, without distinguishing between belief conditions. This result is different from a previous study where the belief of “yes” or “no” nicotine modulated activities in the ventral striatum (but not thalamus) in smokers^10^. We speculate that this discrepancy is mainly due to differences in study design between the current and previous studies. Importantly, the current study design uses e-cigarettes to deliver nicotine to participants were not experienced with vaping, as opposed to the use of traditional cigarettes that smokers were highly experienced with in previous work^10,11^. This “incongruency effect” could have removed conditioned responses related to smoking in the smokers in the current study, as substance-dependent humans are known to be sensitive to subtleties in sensory cues associated with the medium through which the drug is delivered^50,51^. As the striatum is heavily involved in reinforcement learning, it is not surprising that striatal activities showed a response to instructed beliefs in such study design where the mere presence of a cigarette could induce strong conditioned effects. Thus, the identified finding regarding the thalamic circuit represents a mapping between beliefs and neurobiology that is less dependent on conditioned effect as reported in previous work. Furthermore, the average nicotine level was higher in the current study (∼0.8 mg from vaping) than that of our own previous work (∼0.6 mg)^10^. Because the thalamus contains a higher density of nAChRs than the striatum, a higher level of consumed nicotine might amplify thalamus-related activities that are primarily tied to nicotine’s pharmacological effects in this study, as opposed to learned effects in the previous study4,10,11.

In sum, our study provides insight into how a thalamic circuitry represents nicotine-related belief “dosage” in a manner that resembles pharmacological dose-dependent effects. Elucidating the precise mapping mechanism between beliefs and brain states might be important for understanding the key roles cognitive constructs play in human addiction, heterogenous responses to pharmacological treatments^5^, and neural mechanisms of psychotherapeutic effects. As such, this finding opens up new avenues for systematically leveraging the impact of narratives on the brain in mental health research and treatment.

## Materials and Methods

### Participants

The study was approved by the Institutional Review Board of the University of Texas at Dallas and the University of the Texas Southwestern Medical Center where data collection was conducted. All participants signed informed consent before participating in the study.

#### Smokers

Using a similar fMRI learning task and factorial design, a previous study of belief-drug interaction in nicotine addiction (N = 24 per condition) yielded an effect size of Cohen’s d = 0.69 for reward learning. Based on this, we estimated an n = 20 in each belief condition in the final sample would provide 90% power to detect an effect of this magnitude at alpha = 0.05 (two-tailed). Further, sample size calculation with G*Power V3.1.9.7. assuming a three-measurements repeated-measures ANOVA F-test with an effect size of 0.4, alpha = 0.05, and power = 0.95, suggested a minimally required sample size of 18 participants.

Based on this power calculation, we recruited nicotine-dependent adult participants from the Dallas-Fort Worth (DFW) metropolitan area (total n=23 and final n=20). Inclusion criteria include an age of 18 years and older, normal or adjusted to normal vision, and smoking a minimum of 10 cigarettes a day for a period exceeding one year but with no prior experience with vaping devices or current attempt to quit smoking. Exclusion criteria included the use of illicit drugs in the past two months, a history of traumatic brain injury, any current substance abuse (excluding nicotine and alcohol), any contraindication to MRI, or previous or current psychiatric, neurological, or major medical conditions. Twenty-three participants enrolled in this study and underwent three fMRI sessions, spaced about one week apart. Three participants were excluded from analysis for the following reasons: one participant was excluded due to software malfunction, one due to falling asleep in the scanner and one due to loss of behavioral data for one of the scanning sessions. The final sample therefore comprised of 20 participants (6 females, age: 41.1± 11.97 years, age range: 24-61 years). Participants were all right-hand dominant.

#### Non-smoking healthy controls (HC)

In an exploratory analysis, we compared neural activities of the nicotine addicted cohort to those of a healthy controls (HC) cohort which engaged in the same task in the same imaging facility. Thirty-three healthy volunteers (15 females, aged 28 ± 9 years) were recruited for the study using similar criteria as smokers, other than nicotine addiction being an additional exclusion criterion. The sample size for HCs was larger than the required n for smokers as HC data were taken from another study with different overall design and hypotheses.

Two participants were excluded from neural analyses due to excessive head movement (>3 mm), leaving a final sample size of 31.

### Experimental design

Upon arrival at the laboratory, participants completed demographic, mental health (Positive and Negative Affect Schedule, Beck’s Depression Inventory, Empathy Quotient, Toronto Alexithymia Scale, Behavioral Inhibition System and Domain-Specific Risk-Taking questionnaires), general substance and alcohol use (Drug Abuse Screening Test, Short Michigan Alcohol Screening Test), and nicotine-specific surveys (Fagerström Test for Nicotine Dependence, Wisconsin Withdrawal Scale, Shiffman-Jarvik Withdrawal scale). Participants provided saliva samples for measuring cotinine, the primary metabolite of nicotine. Saliva samples were collected using a passive drool method until a volume of 1.8 – 2ml was obtained. They were then coded and stored in designated freezers until sent for analysis. Participants’ exhaled carbon monoxide (CO) levels served as proxy for their satiety status. These were acquired by a Smokerlyzer (coVita micro+basic, Santa Barbara, CA) prior to e-cigarette vaping in each session. The measurement took place in a designated behavioral testing rooms adjacent to the scanners. Participants continuously exhaled through a designated straw until a measurement appeared on screen.

For nicotine delivery, we used the “blu” e-cigarette atomizer (blu, UK) with disposable 1.2% nicotine cartridges in the ‘classic tobacco’ flavor. Following the fMRI scans, participants repeated the state-based series of surveys and provided a second saliva sample.

Three participants’ data were removed from cotinine analysis: two due to cotinine readings exceeding 3 standard deviations from the mean of the cohort and one due to missing data. Data for all three sessions (instructed beliefs conditions) were discarded from this analysis. Three participants’ data (non-overlapping with the previous omission) were removed from CO analysis: two due to readings exceeding 3 standard deviations from times the mean of the cohort and one due to missing data. Once again, data for all three sessions (instructed beliefs conditions) were discarded from this analysis.

Prior to vaping, the e-cigarette cartridge was weighed three times on a high precision scale and the average of the three measurements was logged as the baseline weight of the cartridge. A similar procedure was done post-vaping and the change in cartridge weight represented the amount of nicotine liquid consumed by the participant. Using a double-blind procedure, neither participants nor the research assistants (M.H. and A.K.) responsible for data collection had prior knowledge about the true nicotine content in the e-cigarettes. The order of belief conditions was randomly assigned for each participant. The e-cigarette cartridges were carefully re-labelled as ‘low’, ‘medium’, or ‘high’ by the PI (X.G.) herself to avoid un-blinding by either the participant or the research assistants.

Notably, the same type of cartridges containing 1.2% nicotine were used across all participants and sessions. Research assistants (M.H. and A.K.) who interacted with the participants adhered to a fixed text during the manipulation stating: “The cartridge you will use today will contain: [mild-to-no nicotine] / [a medium amount of nicotine] / [a high amount of nicotine].” These experimenters also made sure that participants used the e-cigarette properly, the device was well powered, and that vapor was visible. Participants were told they could vape as much as they wish for the next 20 minutes and were left alone to vape. After 20 minutes they were questioned about any issues with the e-cigarette. Participants were then prompted to reply how they would rate the strength of the nicotine in the e-cigarette on a scale of 0 to 10, compared to a normal cigarette.

### Cotinine detection in saliva

#### Chemicals and reagents

Optima LC-MS grade acetonitrile, water and methanol were purchased from Fisher Scientific (New Jersey, USA). Reagent grade ammonium formate was purchased from Sigma Aldrich (Missouri, USA). Cotinine was purchased from Sigma Aldrich. Rac-cotinine-d3 was obtained from Toronto Research Chemicals (Ontario, CAN). All other chemicals and reagents were of analytical grade and used without further purification. Blank human saliva procured from primary investigator. (New York, USA).

#### Preparation of stock and working solutions of analyte and internal standard

Primary stock solutions of cotinine for the calibration curve (CC) and quality control samples (QC) were prepared from a 1 mg/mL stock solution in methanol. Stock solutions of cotinine were stored at - 20 °C, and subsequent dilutions were conducted using water. Primary stock solutions of the bioanalytical method’s internal standard (IS) d3-cotinine were prepared by accurately weighing d3-cotinine and dissolving in methanol to yield a 1 mg/mL stock solution. Stock solutions of d3-cotinine were stored at -20 °C, and subsequent dilutions were conducted using water. For spiking of saliva samples with cotinine, a working stock solution of 1000 ng/mL cotinine in water was prepared and stored at -20 °C. For spiking of saliva samples with d3-cotinine, a working stock solution of 300 ng/mL d3-cotinine in water was prepared and stored at -20 °C.

#### Preparation of calibration curve and quality control samples for analysis of saliva samples

Cotinine was validated over a calibration range that supports low concentration and high concentration samples. The calibration curve ranged from 5 ng/mL to 1000 ng/mL. Calibration curve samples were prepared by spiking 1µL of the working stock (1 mg/mL cotinine in water) into 250 µL of saliva. This represented the top calibration curve point (i.e., the upper limit of quantification or ULOQ). The remaining calibration curve samples were prepared by serial dilution of the ULOQ standard in saliva. Quality control was prepared in a similar fashion by spiking 80 µL of the working stock (1000 ng/mL cotinine in water) into 250 µL of plasma. This represented the high-quality control standard (HQC). The medium-quality control standard (MQC) and low-quality control standard (LQC) were prepared by serial dilution of the HQC standard in saliva. Spiking volume of the working standard did not exceed 5% of the matrix volume. QCs for the calibration curve were prepared at 5, 30, and 80 ng/mL.

#### Saliva Collection

Patients provided a saliva sample collected at various time-points throughout a session. Samples were collected using a passive drool method (i.e., Salimetrics ‘SalivaBio’ collection aid). Saliva samples were analyzed for cotinine concentrations using a validated LC-MS/MS method.

#### Saliva Sample Preparation

Acetonitrile (350 µL) and 100 µL (300 ng/mL) d3-cotinine was added to a 250 µL aliquot of saliva. The resultant mixture was centrifuged for 5 min at 5000 rpm at a temperature of 4L. Five hundred microliters (500µL) of the clear supernatant, was removed, placed in a new Eppendorf tube, and dried using a SpeedVac. Samples were reconstituted using 250 µL of water.

#### HPLC operating conditions

A Shimadzu CBM-20A Nexera X2 series LC system (Shimadzu Corporation, Kyoto, Japan) equipped with degasser (DGU-20A) and binary pump (LC-30AD) along with auto-sampler (SIL-30AC) and (CTO-30A) column oven. The autosampler was maintained at 15 °C. An injection volume of 1 µL was used and chromatographic separation was achieved using a Kinetex Biphenyl (2.6 µm, 50 × 2.1mm) column. The mobile phase, consisting of 2mM ammonium formate in water (pump A) and methanol:water (95:5) with 0.2% formic acid (pump B) used for the method. The mobile phase pumped using a gradient program at a flow rate of 0.8 mL/min into the mass spectrometer electrospray ionization chamber in positive polarity. Gradient program initiated with 5% of B and maintained for 1.0min, then ramped to 75%B by 2.5min and maintained at 75%B until 3.5min, changed back to 5%B by 4.0min and maintained until 6.01min at the end system controller stop command.

#### Mass spectrometry operating conditions

Quantitation was achieved employing electrospray ionization in positive ion mode for the analytes using a SCIEX QTRAP 6500+ mass spectrometer (Redwood, CA, USA) equipped with the Turbo V source operated at 550 ºC. The nebulizer gas, auxiliary gas, curtain gas, CAD gas were set at 45, 45, 30 psi and ‘medium’, respectively. The declustering potential (DP), collision energy (CE), entrance potential (EP) and collision cell exit potential (CXP) were 141, 31, 10, 10 V for (Cotinine-1); 141, 47, 10, 8 V for (Cotinine-2); 141, 31, 10, 12 V for (d3-Cotinine-1); and 141, 29, 10, 8 V for (d3-Cotinine-3), respectively. Detection of the ions was carried out in the multiple-reaction monitoring mode (MRM), by monitoring the precursor > product transitions of 177.0 > 80 and 177.0 > 98.0 (sum over 2 MRMs) for cotinine and 181.2 > 80.1 and 181.2> 101.1 (sum over 2 MRMs) for d3-cotinine. The data obtained were processed by Analyst software™ (version 1.6.3).

#### Method Validation

The methods for analysis of cotinine in saliva were validated according to the United States FDA’s May 2018 Guidance for Industry on ‘Bioanalytical Method Validation’. The method was found to have acceptable sensitivity, selectivity, matrix effect, linearity, accuracy, precision, recovery, dilution integrity and stability.

### Value-based decision-making task

This task was developed based on a previous investment task^10,26^ but with the modification that participants were allowed to place both positive (‘invest’) and negative (‘short’) bets. Briefly, participants were allocated an initial sum of 100 monetary units (i.e., their portfolio) at the beginning of the experiment which could be invested in stock markets. Participants were informed that their final payment would be scaled according to their actual gains or losses in the task. Each participant played a total of 10 markets per visit, each consisting of 20 trials. The stock market prices in the task were chosen from true historical stock market prices. Each task block commenced by a caption titled ‘new market’ followed by a graphic display of past market dynamics.

In each trial *t*, the participant observes the price history of a stock market (including the trial before, p_t-1_) and places a bet, *b*_*t*_. Next, a new market price p_t_ is revealed, and portfolio amounts are updated to reflect the recent outcome. The fractional market return *r*_*t*_, is defined as r_t_ = (p_t_ - p_t−1_) / p_t−1_. In each of the 20 trials, participants had unconstrained time to decide on their investment moves. Participants were able to choose to either invest normally (if they think the price will go up) or short sell (if they think the price will go down). Notably, shorting the market would result in gaining from market drops. Thus, people could benefit from either a positive or negative price change and the absolute value of market return |*r*_*t*_| represents the magnitude of the potential gain. Participants provided their choice using a slider bar and finalized their choice by a button press. Following a 750 ms delay, the new market price was revealed and the fractional change in market price was applied to the portfolio. In later analyses this event is termed ‘market reveal’. Each trial concluded in a 750 ms inter-trial interval in which the slider turned gray and became unresponsive. A total of 30 different markets were used across all three visits. Mean session duration in the stock market task was 14.91 ± 3.06 minutes and did not differ across conditions (rmANOVA *F*(2,59) = 0.28, *P* = 0.76).

### Imaging acquisition and preprocessing

Whole-brain anatomical and functional MRI data were acquired on a Philips Achieva scanner with a 3T field strength. High-resolution T1-weighted scans (1.0 × 1.0 × 1.0 mm) were acquired using a 3D magnetization prepared rapid gradient-echo (MPRAGE) sequence. Functional images were acquired using echo-planar imaging (EPI) and tilted 30° from AC-PC axis. The detailed settings for the functional imaging were repetition time (TR) = 2,000 ms, echo time (TE) = 25 ms, flip angle = 90°, voxel size = 3.4 × 3.4 × 4.0 mm, 38 slices. The average number of functional images acquired was 457.37 ± 91.67. All imaging data were preprocessed using standard statistical parametric mapping (SPM12, Wellcome Department of Imaging Neuroscience) algorithms *(fil*.*ion*.*ucl*.*ac*.*uk/spm)*. Functional images were applied a slice time correction.

To account for large head movements often caused by participants’ coughing during the scan, we used the ArtRepair toolbox^52^ to examine and repair volumes with large motion artifacts. We used the *art_motionregress* and *art_global* modules of the single subject pipeline. The ArtRepair algorithm was further used to generate the motion parameters to be included in the GLM design matrix. Volumes were examined for fast head movements using the automated defaults such that volumes with movement of >0.5 mm/TR were tagged and interpolated with the nearest usable volumes. Overall, 6 out of 60 scans were repaired. The mean functional images for each subject were co-registered to the subject’s high-resolution T1 structural scan, using a 12-parameter affine transformation. The participant’s T1 image was segmented into gray and white matter and then normalized using nonlinear basis functions to the Montreal Neurological Institute (MNI) space with the functional images normalized to the template and resampled into 3.4 × 3.4 × 4-mm functional voxels. Functional images were smoothed spatially using a 6 mm full-width at half-maximum Gaussian kernel. A temporal high-pass filter of 128 Hz was applied to the fMRI data, and temporal autocorrelation was modeled using a first-order autoregressive function.

### Statistical analysis

Throughout this study, we used a within-participant repeated measures ANOVA implemented in MATLAB (*anova_rm*) to assess differences between the three conditions of instructed belief. Normality was assessed with Shapiro-Wilk tests wherever appropriate. During analysis of the various controls if data for one of the sessions was missing, that participant was excluded from this specific analysis. For neural data, we specified the statistical thresholds and rationale in the fMRI methods sections below. In the case of between-group comparison between smokers and HC, we used two-sample t-tests, conducted separately for each level of instructed belief.

### Behavioral modeling

We examined the impact of the value signal of market return |r_t_| on participants choice behavior, operationalized as their next bet, |b_t+1_|, using a linear mixed-effects multiple-regression model. The final return of each market was excluded from the regression, as there was no investment decision following the final market segment. Similarly, the first trial was also removed since it had no preceding investment decision. In line with previous investigations^10^, our parameter of interest was instructed belief, expressed as a 3-level interaction (the first level, i.e., ‘low’ belief served as baseline) modulating market return, |r_t_| ^10^.

In order to test whether there was an interacting or a moderating effect of belief on the relationship between market return and next bet, we first tested two plausible models, with- and without an interaction of |*r*_*t*_| and instructed belief. The results suggested that the interaction effect did not improve model fit. We therefore modeled choice behavior as follows:

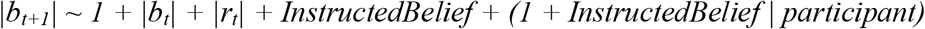

Multiple regression was carried out in R (RStudio 1.1.463, 2018) using the ‘lmer’ function as follows in the ‘lme4’ package *P* values were approximated via Satterthwaite’s degrees of freedom method. The inclusion of instructed belief as a random effect was guided by the notion that the effects of belief are likely heterogenous across the cohort. This intuition was backed up by model comparison between the two options (with / without belief as a random effect) using the ‘anova’ function (*P* < 2.2e-16).

### General linear modeling (GLM) of fMRI data

Event-related analyses of the fMRI data were conducted using SPM12 (Wellcome Department of Imaging Neuroscience). We conducted general linear modeling (GLM) of the functional scans of each participant bey modeling the observed BOLD signals and regressors to identify the relationship between the task events and the hemodynamic response. Regressors related to all visual and motor events were created by convolving a train of delta functions representing the sequence of individual events with the default basis function in SPM12, which consists of a synthetic hemodynamic response function composed of two gamma functions. The GLM included six separate regressors: (1) new market screen display; (2) market history display; (3) all key presses; (4) market price reveal of trial 1; (5) market price reveal of trials 2–19; (6) market price reveal of trial 20. Additionally, six parameters generated during motion correction were entered as covariates. In the GLM, absolute market return |*r*| was entered as a parametric modulator of market reveal of trials 2–19. We carried out linear contrasts of the parameter estimates to identify the effects in each participant.

Statistical maps from all participants were then entered into a second-level group analysis to implement a random-effects statistical model. A within-subject repeated-measures ANOVA model was conducted for the factor of instructed beliefs (‘low’, ‘medium’, ‘high’). Statistical inference was made based on the *F* statistics derived from whole-brain rmANOVA statistical maps. Significant effects were identified at *P < 0*.*05* family-wise error cluster-corrected at a cluster-defining threshold of *P* < 0.005, uncorrected with a cluster size threshold of *k* = 50. We relied in cluster-extend thresholding in our statistical inference in order to allow sufficient sensitivity to detect effects given the experimental sample size ^53^ while implementing thresholds recommended for a balance between Type I and Type II errors ^54^. Maps were rendered using MRIcroGL v1.2.2.

### Thalamic parcellation and ROI analyses

We extracted parameter estimates from bilateral thalamus using an anatomical mask (WFU Pick atlas^28^. Thalamic parcellations were obtained from the Lead-DBS MATLAB toolbox. Each segmented nucleus or region was transformed into the experimental dataset’s functional space using the MarsBaR toolbox. The THOMAS (Thalamus optimized multi-atlas segmentation) atlas^29^ contains 12 non-overlapping nuclei, three of which (vLA, MGN and MTT) were too small to be meaningfully transformed to our functional space and were therefore not used. As before, we modeled BOLD activity tracking of fluctuations in magnitude of market return, |r|, and carried out a group analysis with a within-subject rmANOVA design per region of the thalamic parcellation. To account for multiple comparisons, we applied the false-detection rate (FDR) correction to the extracted ROIs at *q* = 0.05.

### Permutation analysis

We iteratively shuffled labels for instructed beliefs (‘low’, ‘medium’, ‘high’) within each participant while maintaining their original ratio (i.e., one of each per participant). For each viable permutation (i.e., a permutation whose model estimation converged and yielded any significant voxels), a within-subject rmANOVA was carried out, following which a parameter estimate was derived from the same ROI as the original design matrix to generate a surrogate distribution of beta estimates (*N* = 1,000).

### Classification analysis

We decoded instructed belief conditions (‘low’, ‘medium’, ‘high’) from multivoxel spatial patterns data using a rLDA (regularized linear discriminant analysis classifier, ‘fitdiscr’ function in MATLAB)^30,31^. Input data consisted of 20 participants’ first-level GLM maps X 3 conditions along with corresponding belief condition labels, which were used to train the rLDA. A 10-fold cross-validation sample size was used. To test the model’s performance on we iteratively repeated this process with permuted data partitions (*N* = 10,000 for the whole thalamus / NAcc, *N* = 1,000 for thalamic nuclei) per ROI and compared classification accuracy of ground truth data to the surrogate distribution using non-parametric testing.

### Psycho-physiological interaction (PPI) analysis

PPI analysis provides a measure of change in functional connectivity between different brain regions under a specific psychological context^39^. We defined a seed region – the thalamus – as defined by the WFU anatomical atlas and a psychological context (‘market reveal’ – the presentation of the investment’s return). We then conducted a PPI analysis per condition of instructed beliefs and compared those in a within-subject repeated-measures design. The generated PPI model included the PPI term, the physiological regressor, the psychological regressor, and nuisance regressors of six motion parameters.

A 6 mm spherical ROI was defined based on previous investigation of the neural mechanisms of belief-formation in the vmPFC by Rouault and Fleming^40^. Sphere center was set to reflect the coordinate of peak activation (MNI x = -6, y= 52, z = -10). In the follow-up exploratory analysis, the threshold of significance for the group-level rmANOVA from the PPI regressor was set to be *P* < 0.05 FWE cluster-corrected at a cluster-defining threshold of *P* < 0.005, uncorrected.

## Data availability

Data supporting the findings of this study are deposited in: https://osf.io/3hq6s/

## Code availability

The scripts used for data acquisition and analysis are available in: https://osf.io/3hq6s/

Analyses were conducted using open software and toolboxes available online as described in Materials and Methods (SPM: www.fil.ion.ucl.ac.uk/spm/software/spm12;

R Studio: https://www.rstudio.com/products/rstudio/download/#download;

Lead-DBS: https://www.lead-dbs.org/download/;

MRIcroGL: https://www.nitrc.org/projects/mricrogl/)

## Acknowledgements

We thank staff members at the UT Southwestern Imaging Center for their assistance with scanning, Jennifer Jung and Mark Labinski for their help with developing the fMRI task, and Jessica Maclin for her help with participant recruitment. We also thank Dr. Laura Berner for her advice on the statistical analysis, and Drs. Daniela Schiller and Paul Kenny for their helpful discussions and comments on an earlier version of this manuscript.

## Author contributions

Conceptualization: XG.

Methodology: OP, AS, NB, WCP, XG

Investigation: MH, SN, AK

Visualization: OP

Funding acquisition: XG

Project administration: MH, XG

Supervision: XG

Writing – original draft: OP, WCP, VGF, XG

Writing – review and editing: OP, VGF, XG

## Competing interests

The authors declare that they have no competing interests.

## Funding

National Institute on Drug Abuse grant R01DA043695 (XG)

National Institute on Drug Abuse grant R21DA049243 (XG)

University of Texas, Dallas internal funding (XG)

Mental Illness Research, Education, and Clinical Center at the James J. Peter Veterans Affairs

Medical Center grant MIRECC VISN 2 (VGF)

## Supplementary information

Supplementary Figs. 1 to 8

Supplementary Table 1

## Notes

https://osf.io/3hq6s/

### Competing Interest Statement

The authors have declared no competing interest.

### Summary of Updates

Reformatting from brief to a full research article.

## References

1. Posner, M. I., Petersen, S. E., Fox, P. T. & Raichle, M. E. Localization of cognitive operations in the human brain. Science 240, 1627–1631 (1988).

2. Gu, X., FitzGerald, T. H. B. & Friston, K. J. Modeling subjective belief states in computational psychiatry: interoceptive inference as a candidate framework. Psychopharmacology (Berl). 236, 2405–2412 (2019).

3. Goldstein, R. Z. et al. The Neurocircuitry of Impaired Insight in Drug Addiction. Trends Cogn. Sci. 13, 372–380 (2009).

4. Volkow, N. D. & Baler, R. Beliefs modulate the effects of drugs on the human brain. Proc. Natl. Acad. Sci. U. S. A. 112, 2301–2302 (2015).

5. Kirsch, I. Response expectancy as a determinant of experience and behavior. Am. Psychol. 40, 1189–1202 (1985).

6. Mayberg, H. S. et al. The functional neuroanatomy of the placebo effect. Am. J. Psychiatry 159, 728–737 (2002).

7. Wager, T. D. et al. Placebo-induced changes in FMRI in the anticipation and experience of pain. Science 303, 1162–1167 (2004).

8. Benedetti, F., Mayberg, H. S., Wager, T. D., Stohler, C. S. & Zubieta, J. K. Neurobiological mechanisms of the placebo effect. J. Neurosci. 25, 10390–10402 (2005).

9. Price, D. D., Finniss, D. G. & Benedetti, F. A comprehensive review of the placebo effect: Recent advances and current thought. Annu. Rev. Psychol. 59, 565–590 (2008).

10. Gu, X. et al. Belief about nicotine selectively modulates value and reward prediction error signals in smokers. Proc. Natl. Acad. Sci. U. S. A. 112, 2539–2544 (2015).

11. Gu, X. et al. Belief about nicotine modulates subjective craving and insula activity in deprived smokers. Front. Psychiatry 7, 1–11 (2016).

12. Benowitz, N. L., Jacob, P. & Herrera, B. Nicotine intake and dose response when smoking reduced-nicotine content cigarettes. Clin. Pharmacol. Ther. 80, 703–714 (2006).

13. Bisby, J. A., Leitz, J. R., Morgan, C. J. A. & Curran, H. V. Decreases in recollective experience following acute alcohol: A dose-response study. Psychopharmacology (Berl). 208, 67–74 (2010).

14. Curran, V. H., Brignell, C., Fletcher, S., Middleton, P. & Henry, J. Cognitive and subjective dose-response effects of acute oral Δ9-tetrahydrocannabinol (THC) in infrequent cannabis users. Psychopharmacology (Berl). 164, 61–70 (2002).

15. Watkins, S. S., Koob, G. F. & Markou, A. Neural mechanisms underlying nicotine addiction: Acute positive reinforcement and withdrawal. Nicotine Tob. Res. 2, 19–37 (2000).

16. Huang, A. S., Mitchell, J. A., Haber, S. N., Alia-Klein, N. & Goldstein, R. Z. The thalamus in drug addiction: From rodents to humans. Philos. Trans. R. Soc. B Biol. Sci. 373, (2018).

17. Haber, S. N. & Knutson, B. The reward circuit: Linking primate anatomy and human imaging. Neuropsychopharmacology 35, 4–26 (2010).

18. Shohamy, D. Learning and motivation in the human striatum. Curr. Opin. Neurobiol. 21, 408–414 (2011).

19. Spurden, D. P. et al. Nicotinic receptor distribution in the human thalamus: Autoradiographical localization of [3H]nicotine and [125I]α-bungarotoxin binding. J. Chem. Neuroanat. 13, 105–113 (1997).

20. Paterson, D. & Nordberg, A. Neuronal nicotinic receptors in the human brain. Prog. Neurobiol. 61, 75–111 (2000).

21. Gotti, C. et al. Nicotinic acetylcholine receptors in the mesolimbic pathway: Primary role of ventral tegmental area α6β2* receptors in mediating systemic nicotine effects on dopamine release, locomotion, and reinforcement. J. Neurosci. 30, 5311–5325 (2010).

22. Brody, A. L. et al. Cigarette smoking saturates brain alpha 4 beta 2 nicotinic acetylcholine receptors. Arch. Gen. Psychiatry 63, 907–915 (2006).

23. Brody, A. L. et al. Brain nicotinic acetylcholine receptor occupancy: Effect of smoking a denicotinized cigarette. International Journal of Neuropsychopharmacology vol. 12 305–316 (2009).

24. Brody, A. L. et al. Effect of secondhand smoke on occupancy of nicotinic acetylcholine receptors in brain. Arch. Gen. Psychiatry 68, 953–960 (2011).

25. Petersen, G. O., Leite, C. E., Chatkin, J. M. & Thiesen, F. V. Cotinine as a biomarker of tobacco exposure: Development of a HPLC method and comparison of matrices. J. Sep. Sci. 33, 516–521 (2010).

26. Lohrenz, T., McCabe, K., Camerer, C. F. & Montague, P. R. Neural signature of fictive learning signals in a sequential investment task. Proc. Natl. Acad. Sci. U. S. A. 104, 9493–9498 (2007).

27. Chiu, P. H., Lohrenz, T. M. & Montague, P. R. Smokers’ brains compute, but ignore, a fictive error signal in a sequential investment task. Nat. Neurosci. 11, 514–520 (2008).

28. Maldjian, J. A., Laurienti, P. J., Kraft, R. A. & Burdette, J. H. An automated method for neuroanatomic and cytoarchitectonic atlas-based interrogation of fMRI data sets. Neuroimage 19, 1233–1239 (2003).

29. Su, J. H. et al. Thalamus Optimized Multi Atlas Segmentation (THOMAS): fast, fully automated segmentation of thalamic nuclei from structural MRI. Neuroimage 194, 272–282 (2019).

30. Zander, T. O., Kothe, C., Jatzev, S. & Gaertner, M. Enhancing Human-Computer Interaction with Input from Active and Passive Brain-Computer Interfaces BT - Brain-Computer Interfaces: Applying our Minds to Human-Computer Interaction. in (eds. Tan, D. S. & Nijholt, A.) 181–199 (Springer London, 2010). doi:10.1007/978-1-84996-272-8_11.

31. Mandelkow, H., De Zwart, J. A. & Duyn, J. H. Linear discriminant analysis achieves high classification accuracy for the BOLD fMRI response to naturalistic movie stimuli. Front. Hum. Neurosci. 10, 1–12 (2016).

32. Volkow, N. D., Wang, G. J., Fowler, J. S., Tomasi, D. & Telang, F. Addiction: Beyond dopamine reward circuitry. Proc. Natl. Acad. Sci. U. S. A. 108, 15037–15042 (2011).

33. Mikl, M. et al. Effects of spatial smoothing on fMRI group inferences. Magn. Reson. Imaging 26, 490–503 (2008).

34. Shepherd, G. M. G. & Yamawaki, N. Untangling the cortico-thalamo-cortical loop: cellular pieces of a knotty circuit puzzle. Nat. Rev. Neurosci. 22, 389–406 (2021).

35. Schuck, N. W. & Niv, Y. Sequential replay of nonspatial task states in the human hippocampus. Science 364, (2019).

36. Rushworth, M. F. S. & Behrens, T. E. J. Choice, uncertainty and value in prefrontal and cingulate cortex. Nat. Neurosci. 11, 389–397 (2008).

37. de Kloet, S. F. et al. Bi-directional regulation of cognitive control by distinct prefrontal cortical output neurons to thalamus and striatum. Nat. Commun. 12, 1994 (2021).

38. Steward, T. et al. A thalamo-centric neural signature for restructuring negative self-beliefs. Mol. Psychiatry 27, 1611–1617 (2022).

39. Friston, K. J. et al. Psychophysiological and Modulatory Interactions in Neuroimaging. Neuroimage 6, 218–229 (1997).

40. Rouault, M. & Fleming, S. M. Formation of global self-beliefs in the human brain. Proc. Natl. Acad. Sci. U. S. A. 117, 27268–27276 (2020).

41. Niv, Y. Learning task-state representations. Nat. Neurosci. 22, 1544–1553 (2019).

42. Adem, A. et al. Distribution of nicotinic receptors in human thalamus as visualized by 3H-nicotine and 3H-acetylcholine receptor autoradiography. J. Neural Transm. 73, 77–83 (1988).

43. Wong, D. F. et al. PET imaging of high-affinity a4b2 nicotinic acetylcholine receptors in humans with 18F-AZAN, a radioligand with optimal brain kinetics. J. Nucl. Med. 54, 1308–1314 (2013).

44. Lawrence, N. S., Ross, T. J. & Stein, E. A. Cognitive mechanisms of nicotine on visual attention. Neuron 36, 539–548 (2002).

45. Stein, E. A. et al. Nicotine-Induced Limbic Cortical Activation in the Human BrainL: A Functional MRI Study. Am. J. Psychiatry 155, 1009–1015 (1998).

46. Kumari, V. et al. Cognitive effects of nicotine in humans: An fMRI study. Neuroimage 19, 1002–1013 (2003).

47. Levy, D. J. & Glimcher, P. W. The root of all value: A neural common currency for choice. Curr. Opin. Neurobiol. 22, 1027–1038 (2012).

48. Schuck, N. W., Cai, M. B., Wilson, R. C. & Niv, Y. Human Orbitofrontal Cortex Represents a Cognitive Map of State Space. Neuron 91, 1402–1412 (2016).

49. Barbalat, G., Bazargani, N. & Blakemore, S.-J. The Influence of Prior Expectations on Emotional Face Perception in Adolescence. Cereb. Cortex 23, 1542–1551 (2012).

50. Perkins, K. A. et al. Effects of central and peripheral nicotinic blockade on human nicotine discrimination. Psychopharmacology (Berl). 142, 158–164 (1999).

51. Perkins, K. A., Herb, T. & Karelitz, J. L. Discrimination of nicotine content in electronic cigarettes. Addict. Behav. 91, 106–111 (2019).

52. Mazaika, Whitfield-Gabrieli & Reiss…, A. Artifact repair for fMRI data from high motion clinical subjects. Annu. Meet. … 321 (2007).

53. Woo, C.-W., Krishnan, A. & Wager, T. D. Cluster-extent based thresholding in fMRI analyses: Pitfalls and recommendations. Neuroimage 91, 412–419 (2014).

54. Lieberman, M. D. & Cunningham, W. A. Type I and Type II error concerns in fMRI research: re-balancing the scale. Soc. Cogn. Affect. Neurosci. 4, 423–428 (2009).

