## Supplementary Information for "A thalamic circuit represents dose-like responses induced by nicotine-related beliefs in human smokers"

**This PDF file includes:**

Supplementary Table 1

Supplementary Figs. 1 to 8

**Supplementary Information:**

**Localization of observed effects of belief on thalamic activity**

In an exploratory analysis, we delineated the how beliefs modulated activities in different subnuclei within the thalamus. We used THOMAS – an anatomy-based segmentation to test for effects of belief through ROI analyses on segmented thalamic nuclei^1^. For simplicity we focused on the left thalamus where the overall belief effect was more robust. Given the resolution of our functional scans, we extracted the nine largest ROIs from the atlas.

This finer parcellation of the thalamus identified that several ventral posterior nuclei – notably the centromedian (CM) and lateral geniculate nuclei (LGN), were the primary thalamic nuclei whose reward-tracking activity was modulated by instructed beliefs (rmANOVA, FDR corrected at *q* = 0.05; VPL, Pulvinar, LGN, CM all *P* < 0.05) (**Supp Fig. 2)**.

**Effects of instructed belief on choice behavior in the Stock Market task**.
Regressing participants’ magnitude of next bet, |*b_t+1_|*, uncovered an expected strong positive relationship with the magnitude of the current bet (coefficient: 0.250 ± 0.024, t(1.96) = 10.164, *P* = 2.88E-09), but only a weak contribution by magnitude of market return, *|r_t_|* (coefficient: 0.034 ± 0.009, t(1.07E04) = 0.253, *P* = 0.011). Model intercept was modestly reduced when contrasting the ‘low’ and ‘high’ belief conditions coefficient: -0.058 ± 0.026, t(18.90) = -2.31, *P* = 0.046) (see **Supp Fig. 6** and **Supplementary Table 1** for full statistical information).

**Effects of belief about nicotine on difference learning signals in the brain**

Reward prediction errors derived from differences between expected and actually experienced rewards (temporal difference (TD) errors) are also known to impact choice behavior^2^. The neural correlates of these learning signals were previously localized to key mesolimbic dopaminergic regions such as the striatum. Parameter estimates derived from the anatomical demarcation of the thalamus were not modulated by instructed beliefs (*P* = 0.869). We extracted neural signals associated with TD across the cohort (i.e., conditions pooled, rather than contrasted) in a whole-brain contrast and observed pronounced activation in the nucleus accumbens that tracked TD fluctuations (peak at MNI x = -15, y = 8, z = -14; rmANOVA, *P* < 0.05, FWE whole-brain corrected, **Supplementary Fig. 8a**). However, in line with the negligible modulatory effects of instructed beliefs on overall striatal value-tracking, parameter estimates from NAcc were also unchanged by instructed beliefs (rmANOVA, both *P* > 0.73, **Supplementary Fig. 8b**).

**No difference in motor action in choice behavior across belief conditions**

We compared the average number of button presses and their frequency (presses per second within the input period of the trial across conditions; Mean number of button presses per participant in the ‘high’ instructed beliefs condition trended to be lower than the other two conditions (rmANOVA F(2,38) = 2.82, *P* = 0.072). We also accounted for the actual temporal sequences of the button presses (the bet input was self-paced, therefore the same number of button presses can potentially drive different BOLD signals if pressed rapidly over a short period of time, or more gradually). The frequency of button presses did not differ significantly across belief conditions (frequency of presses within rmANOVA *F*(2,38) = 0.99, *P* = 0.38).

**Supplementary Figures**

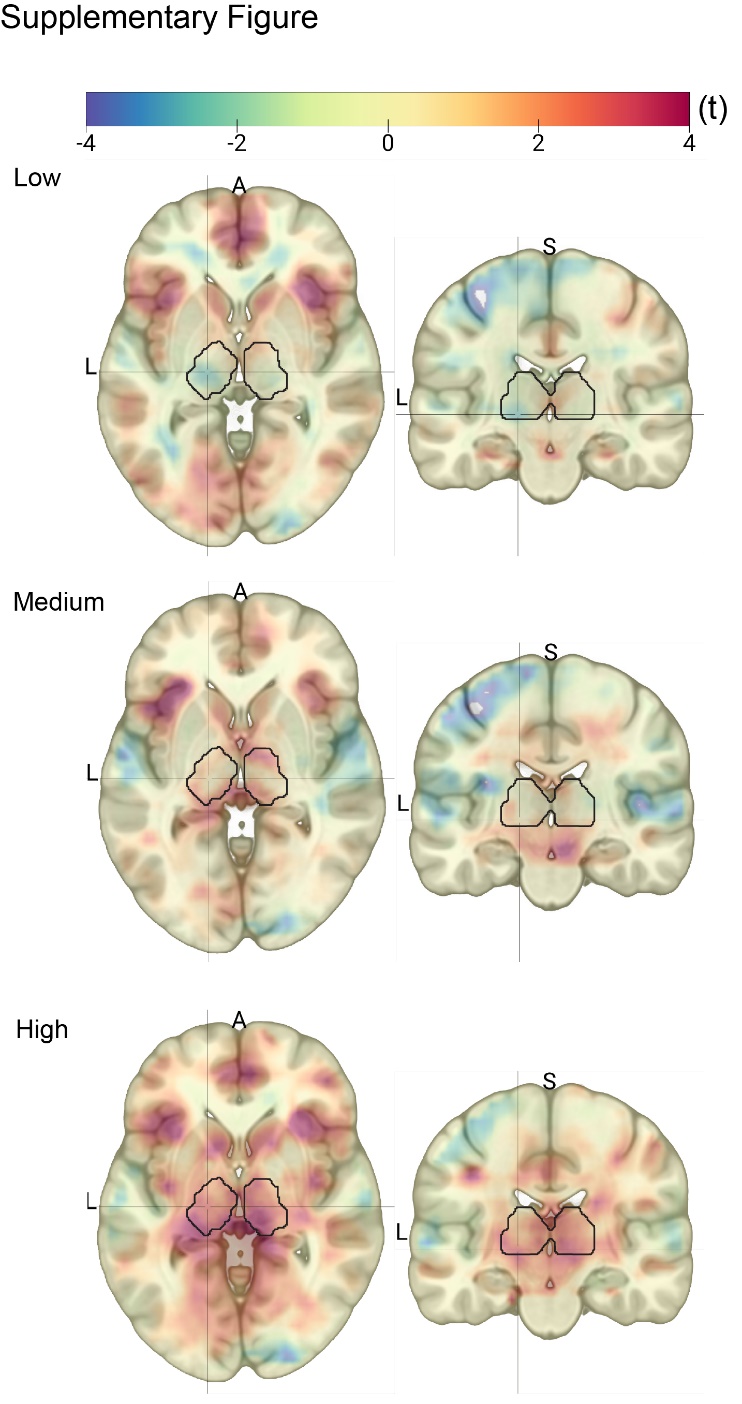

**Supplementary Figure 1**

Whole-brain effects of instructed beliefs about nicotine on value-tracking signals. Each panel portrays the statistical map generated for a single instructed belief condition – ‘low’, ’medium’ and ‘high’. The maps are centered on coordinates of statistical peak at left posterior thalamus detailed in the main text (MNI coordinates: x = -15, y = -19, z = -1). Black contour marks the thalamus. Note that all maps use a common color scale to represent statistical value *(t)* to aid in visualization of changes across conditions*.*

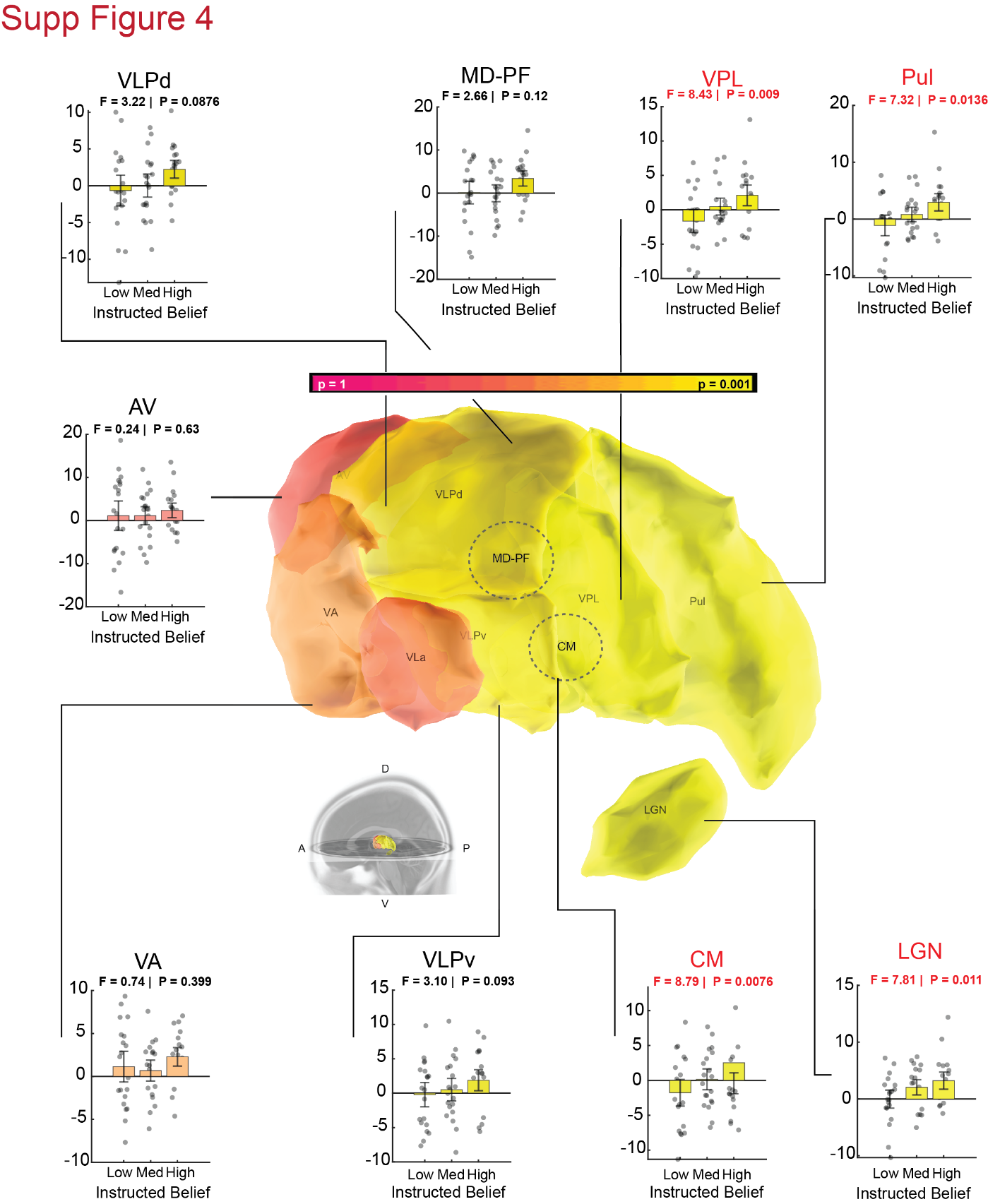

**Supplementary Figure 2**

Center: a color-coded 3D rendering of the segmented left thalamus. Orientation is provided in bottom left inset. Colors denote significance of a repeated-measures ANOVA run on parameter estimates as a function of instructed beliefs. Magenta to yellow color gradient denotes values ranging between *P* = 1 and *P* = 0.001. Each subplot is a bar graph showing parameter estimates extracted from a thalamic subregion for the magnitude of market return, |*r_t_|* per condition of instructed beliefs. Points are participants, jittered horizontally. Values were subjected to per-ROI rmANOVA whose output is displayed in the subplots’ labels. Error bars are SEM. *P* values were then corrected for multiple comparisons with FDR at *q* < 0.05. The red labels denote significant effects of instructed beliefs on thalamic activity. Note the effects of belief were localized to the posterior regions of the thalamus.

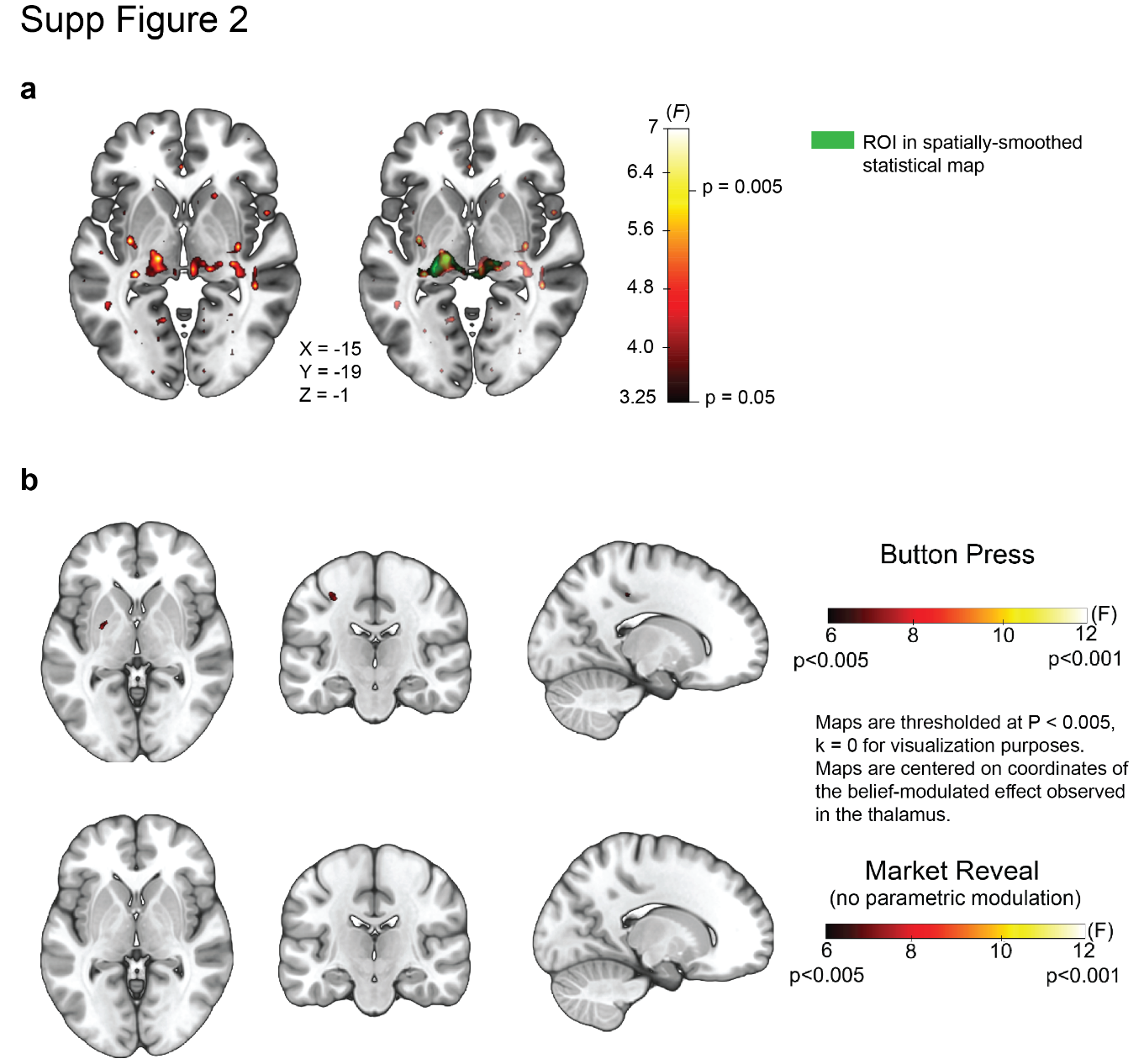

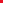

**Supplementary Figure 3**

**(a)** Effects of spatial smoothing on the localization of belief-mediated effects of value tracking in ventral-posterior thalamus **Left:** A whole-brain group analysis depicting effects of instructed beliefs about nicotine. This map displays the same data as shown in Figure 2, but without spatial smoothing applied in the imaging preprocessing pipeline. The event of market reveal, parametrically modulated with the magnitude of market return *|r_t_|* was contrasted between the three conditions of instructed beliefs (‘low’ / ‘medium’ / ‘high’) in a repeated-measures design. The map is centered on coordinates of statistical peak at left posterior thalamus detailed in the main text (MNI coordinates: x = -15, y = -19, z = -1). Heatmap signifies *F* values, corresponding range of *P* values is overlaid. **Right:** same as Left but with the result of the spatially smoothed GLM contrast overlaid in green on top of the non-smoothed statistical map.

Note the smoothed and non-smoothed maps overlap.

**(b)** Instructed beliefs conditions did not differ in activation due to motor outputs and non-parametrically modulated visual cues. **Top:** A whole-brain group analysis depicting effects of instructed beliefs about nicotine. Neural activity associated with events of button press was contrasted between the three conditions of instructed beliefs (‘low’ / ‘medium’ / ‘high’) in a repeated-measures design. Statistical thresholds were set to be comparable with the main effect at rmANOVA, *P* < 0.005, see Fig. 2. The map is centered on coordinates of statistical peak at left posterior thalamus detailed in the main text (MNI coordinates: x = -15, y = -19, z = -1). Heatmap signifies F values, corresponding range of *P* values is overlaid. **Bottom:** same as Top, but for events of market-reveal without parameter modulation by market return, essentially testing for belief-modulated thalamic response for the onset of monetary outcome as visual cue, devoid of value.
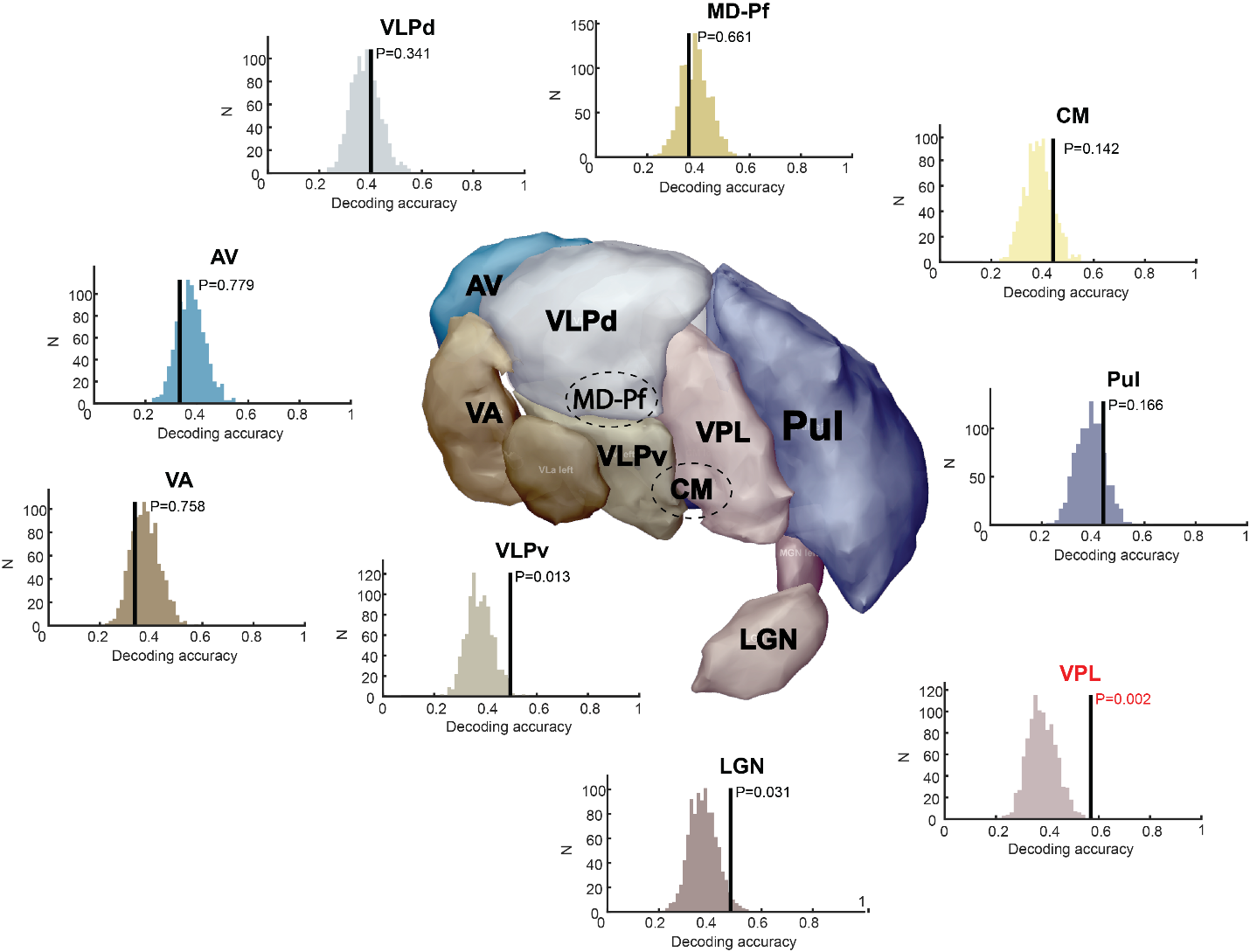

**Supplementary Figure 4**

Center: a 3D rendering of the segmented left thalamus. Each subplot is a histogram comprised of decoding accuracy for the neural data with shuffled labels. Vertical black line denotes decoding accuracy for ground truth data. Histograms are surrogate distributions comprised of decoding accuracy for the same neural data with shuffled labels. Histogram colors are arbitrary. P value is derived non-parametrically through permutation tests and corrected for multiple comparisons with FDR at *q* < 0.05 (colored red).

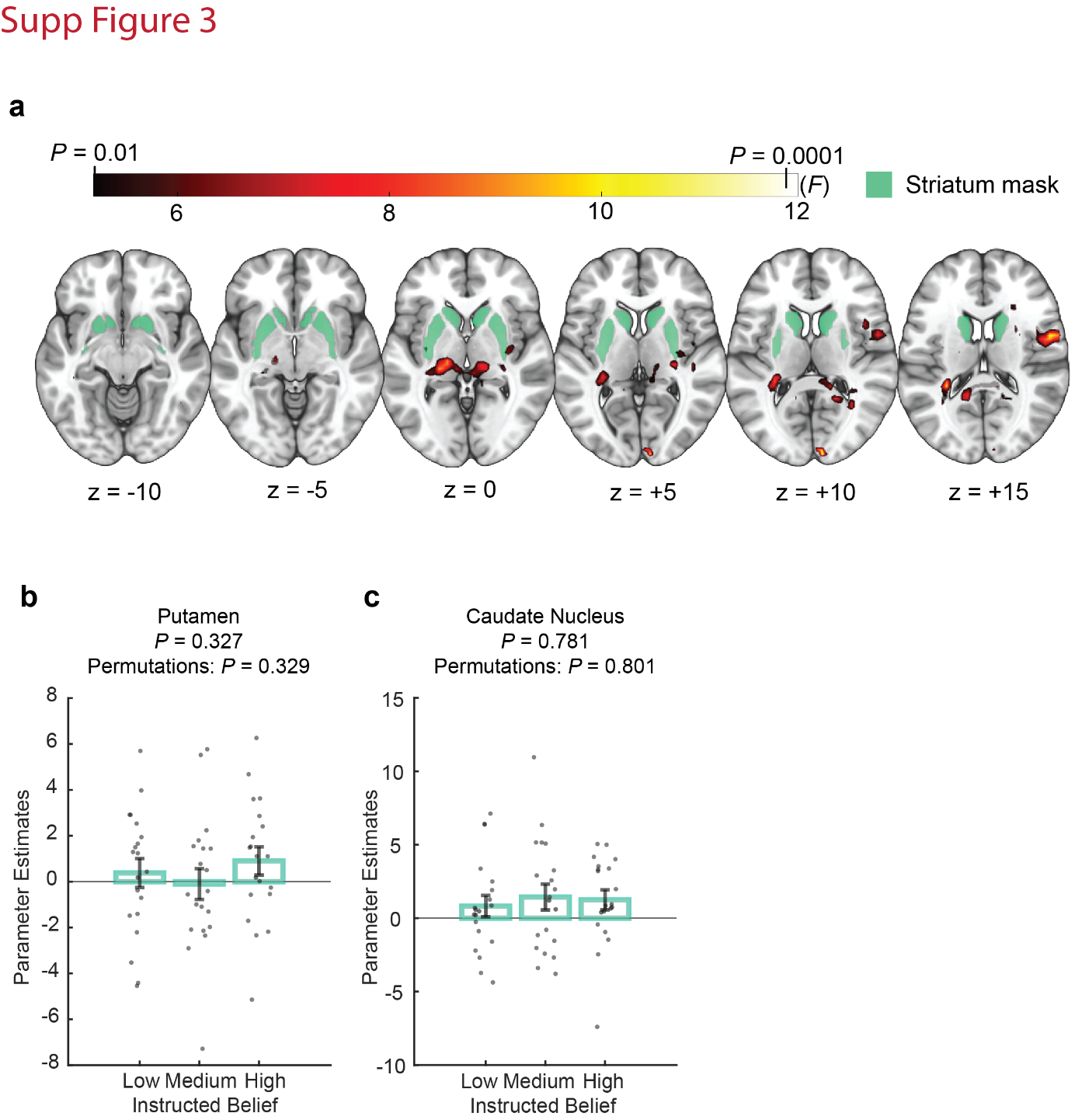

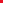

**Supplementary Figure 5**

**(a)** A whole-brain group analysis depicting effects of instructed beliefs about nicotine. The event of market reveal, parametrically modulated with the magnitude of market return *|r_t_|* was contrasted between the three conditions of instructed beliefs (‘low’ / ‘medium’ / ‘high’) in a repeated-measures design. A parcellation of the major striatal regions is highlighted in purple (caudate, putamen, nucleus accumbens). The five axial planes provide coverage of the striatum ranging from z = -5 to z = +15 in MNI space. Heatmap signifies F values, corresponding range of *P* values is overlaid.

(**b**) Region-of-interest analysis on the effect of belief about nicotine on putamen activity. Market reveal, parametrically modulated with magnitude of market return |r_t_| was contrasted between conditions of instructed beliefs (‘low’ / ‘medium’ / ‘high’) in a repeated measures complemented with a non-parametric condition permutation test. Bars depict group means per condition of instructed beliefs. points are participants, jittered horizontally. Error bars are SEM.

(**c**) same as (**b**), but for the caudate nucleus.

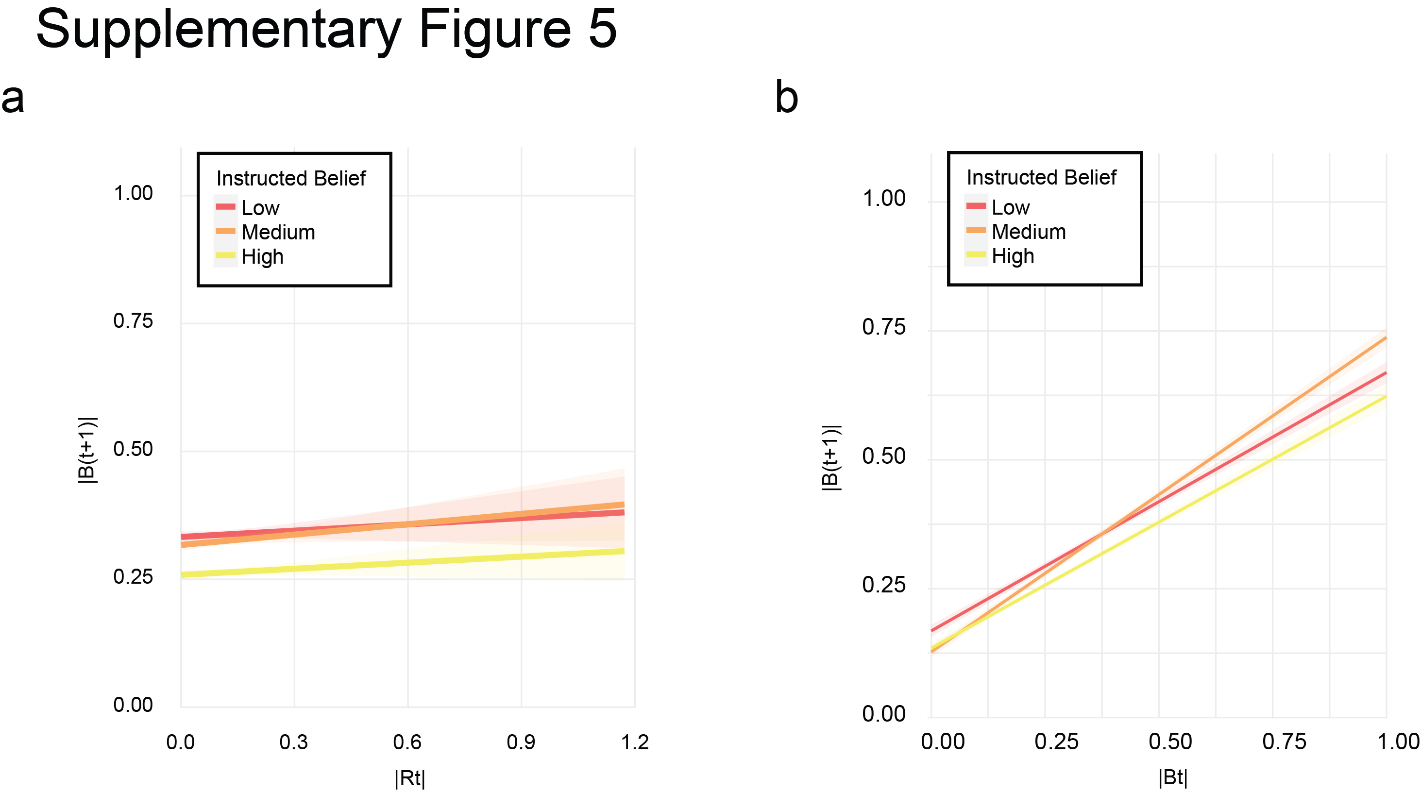

**Supplementary Figure 6**

(a) Choice behavior in the stock market task. Linear modeling of the relationship between magnitude of market return at time (t) (|Rt|), and magnitude of bet size in the next trial (|Bt+1|), as a function of instructed belief about nicotine strength. Data are collapsed across all trials and participants. Shaded area denotes standard error bounds.
(b) Same as **(a)** but for relationship between magnitude of bet size in time (t) (|Bt|), and magnitude of bet size in the next trial (|Bt+1|).

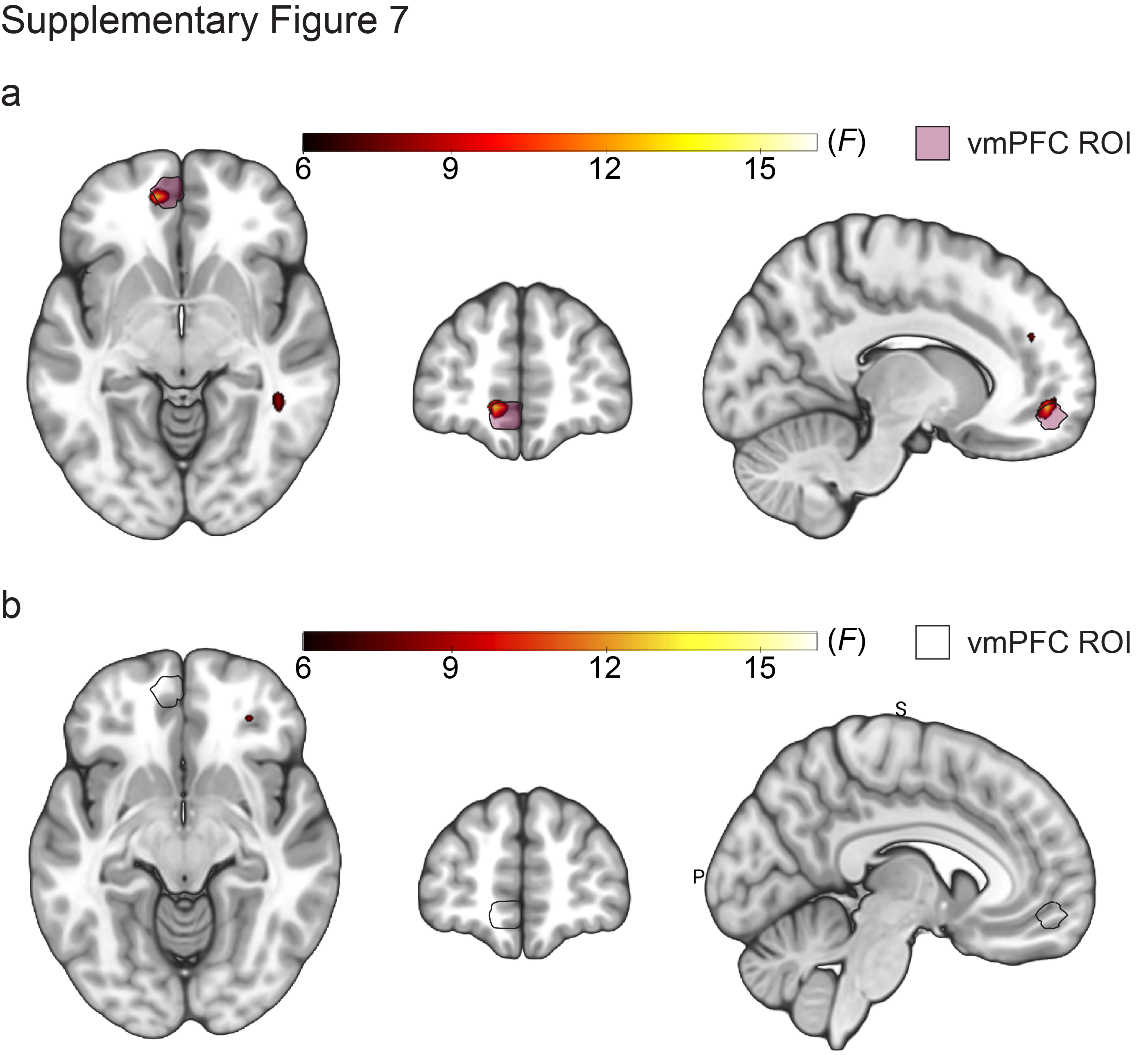

**Supplementary Figure 7**

(a) Location of spherical ROI located in the vmPFC (pink), used to investigate effects of instructed beliefs on the psychophysiological interaction (PPI) between the thalamus and the vmPFC. Visualization is centered at the coordinate of peak activation (MNI x = -10, y = 50, z = -7) based on a previous investigation of the neural mechanisms of belief-formation in the vmPFC.
(b) Same as **(a)** but for PPI analysis using bilateral nucleus accumbens as seed. Visualization is centered at the same coordinates of vmPFC activation reported in **(a)**.

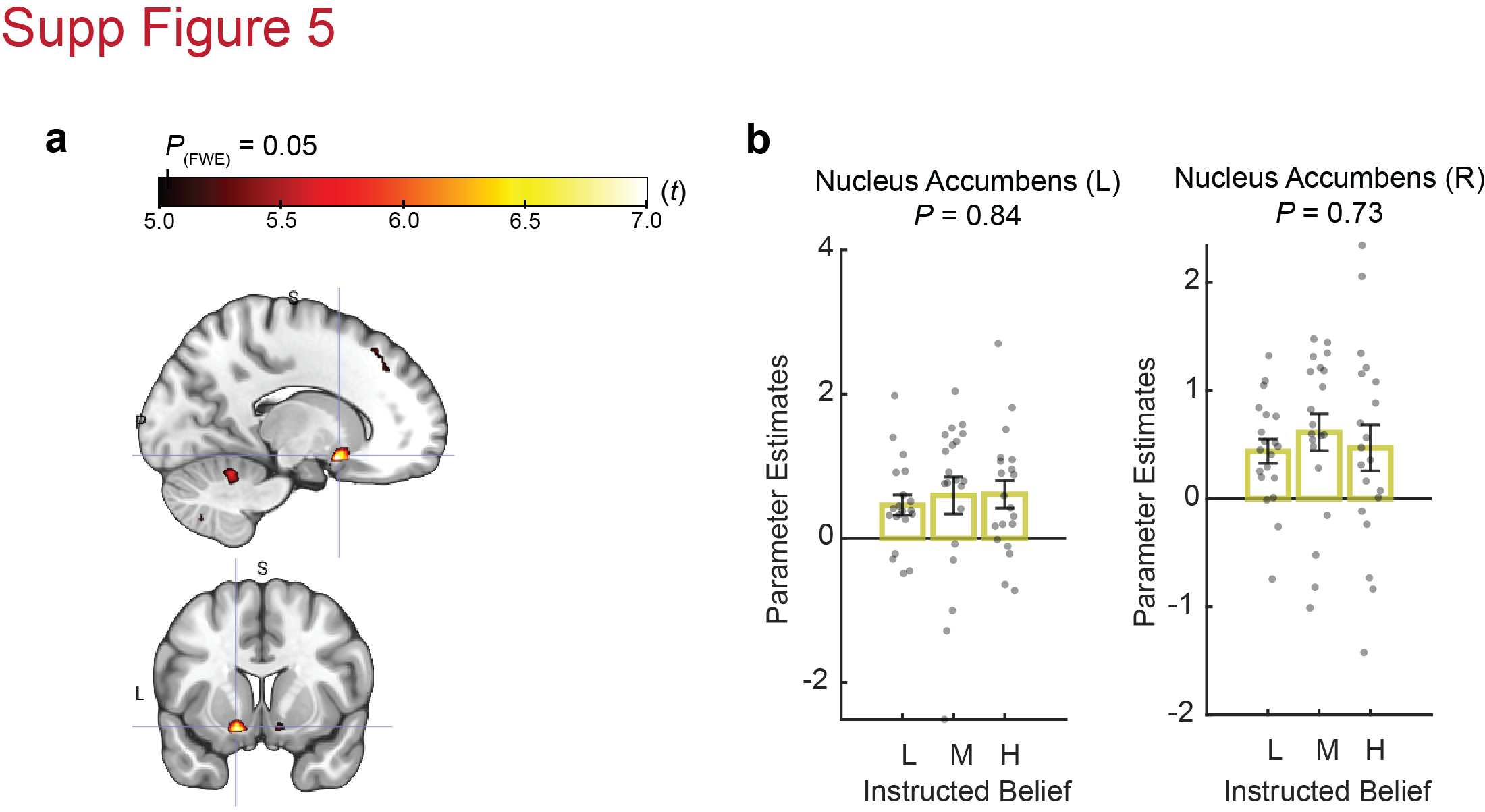

**Supplementary Figure 8**

(**a**) Belief about nicotine did not modulate temporal difference learning signals in ventral striatum. A whole-brain group analysis statistical map depicting brain response to the event of market reveal parametrically modulated with per-trial temporal difference (TD) signals (the actual monetary gain minus the expected gain). Note that this map is derived from statistical maps pooled across all conditions of instructed beliefs. Crosshair is centered on left nucleus accumbens (MNI coordinates: x = -15, y = 8, z = -14). Heatmap signifies *t* values, corresponding cutoff in *P* value FWE-corrected at *P* < 0.05 is overlaid.

(**b**) Parameter estimates extracted from left and right nucleus accumbens (left and right panels, respectively) for temporal difference (TD) signals. Bars depict group means per condition of instructed beliefs. points are participants, jittered horizontally. Error bars are SEM. Note that despite being consistently high, parameter estimates for TD signals suggest they were not affected by instructed beliefs in the nucleus accumbens (both *P* > 0.73). L/M/H labels signify ‘low’, ‘medium’, and ‘high’ states of instructed beliefs about nicotine strength.

**Supplementary Tables:**

| **Fixed effects** | **Estimate** | **SD** | **df** | **t** | ***P*** |
| --- | --- | --- | --- | --- | --- |
| Intercept | 2.51E-01 | 2.46E-02 | 1.97E+01 | 10.164 | 2.88E-09 |
| \|B_(t)_\| | 2.46E-01 | 9.36E-03 | 1.08E+04 | 26.258 | <2.00E-16 |
| \|R_(t)_\| | 3.48E-02 | 1.37E-02 | 1.07E+04 | 2.536 | 0.0112 |
| Instructed belief (low vs medium) | -9.31E-03 | 2.79E-02 | 1.89E+01 | -0.334 | 0.742 |
| Instructed belief (low vs high) | -5.58E-02 | 2.62E-02 | 1.89E+01 | -2.131 | 0.0464 |

**Supplementary Table 1**

Choice behavior – betting in the Stock Market task. A general linear model was applied separately per participant. Presented here are averages across all 20 smoker participants. b(t), market bet at time t. r(t), market return at time t. SD, standard deviation. SE, standard error, df, degrees of freedom.

**References:**

1.	Su, J. H. *et al.* Thalamus Optimized Multi Atlas Segmentation (THOMAS): fast, fully automated segmentation of thalamic nuclei from structural MRI. *Neuroimage* **194**, 272–282 (2019).

2.	Schultz, W., Dayan, P. & Montague, P. R. A Neural Substrate of Prediction and Reward. *Science (80-. ).* **275**, 1593–1599 (1997).
